# Genesis and regulation of C-terminal cyclic imides from protein damage

**DOI:** 10.1101/2024.08.09.606997

**Authors:** Wenqing Xu, Zhenguang Zhao, Matthew Su, Atul Jain, Hannah C. Lloyd, Ethan Yang Feng, Nick Cox, Christina M. Woo

**Author notes:** **Corresponding author:** Christina M. Woo **Email:**.

## Abstract

C-Terminal cyclic imides are post-translational modifications (PTMs) that can arise from spontaneous intramolecular cleavage of asparagine or glutamine residues resulting in a form of irreversible protein damage. These protein damage events are recognized and removed by the E3 ligase substrate adapter cereblon (CRBN), indicating that these aging-related modifications may require cellular quality control mechanisms to prevent deleterious effects. However, the factors that determine protein or peptide susceptibility to C-terminal cyclic imide formation or their effect on protein stability have not been explored in detail. Here, we characterize the primary and secondary structures of peptides and proteins that promote intrinsic formation of C-terminal cyclic imides in comparison to deamidation, a related form of protein damage. Extrinsic effects from solution properties and stressors on the cellular proteome additionally promote C-terminal cyclic imide formation on proteins like glutathione synthetase (GSS) that are susceptible to aggregation if the protein damage products are not removed by CRBN. This systematic investigation provides insight to the regions of the proteome that are prone to these unexpectedly frequent modifications, the effects of this form of protein damage on protein stability, and the biological role of CRBN.

## Introduction

Biological systems and their proteins are subject to unavoidable protein damage events, resulting in post-translational modifications (PTMs) that can disrupt protein structure, stability, and function in the absence of protein repair and removal processes(1). While these non-enzymatic protein damage events can arise from intermolecular reactions between metabolites and the protein, such as oxidation(2), glycation(3) and nitration(4), many other protein damage events occur via intramolecular processes intrinsic to the protein’s primary or secondary structure, such as autoproteolysis(5), isomerization(6, 7), racemization(8, 9), elimination(10, 11) and deamidation at asparagine(6, 12). These forms of protein damage emerge spontaneously over time, reflecting the molecular process of aging.

Asparagine residues can also undergo an alternate intrinsic protein damage event resulting in the C-terminal cyclic imide modification. C-terminal cyclic imide formation is an irreversible event that arises from the nucleophilic addition of the side chain amide to the adjacent backbone carbonyl, leading to cleavage of the protein that affords a N-terminal fragment bearing the C-terminal cyclic imide modification and a C-terminal fragment (**Fig. 1A**). This relatively underappreciated chemical mechanism has been observed to form primarily from cyclization of asparagine (cN) and to a lesser extent on glutamine (cQ) in peptide and protein literature and mass spectrometry experiments(6, 13–17). Additionally, C-terminal cyclic imides have been recently annotated in biological systems on long-lived proteins, such as crystallins and aquaporin 0 in the eye lens, which may drive crosslinking events associated with cataracts(18–20). If unmitigated, protein damage from these cleavage events and their hydrolysis products may accumulate and produce undesirable effects in biological systems.

**Figure 1.**
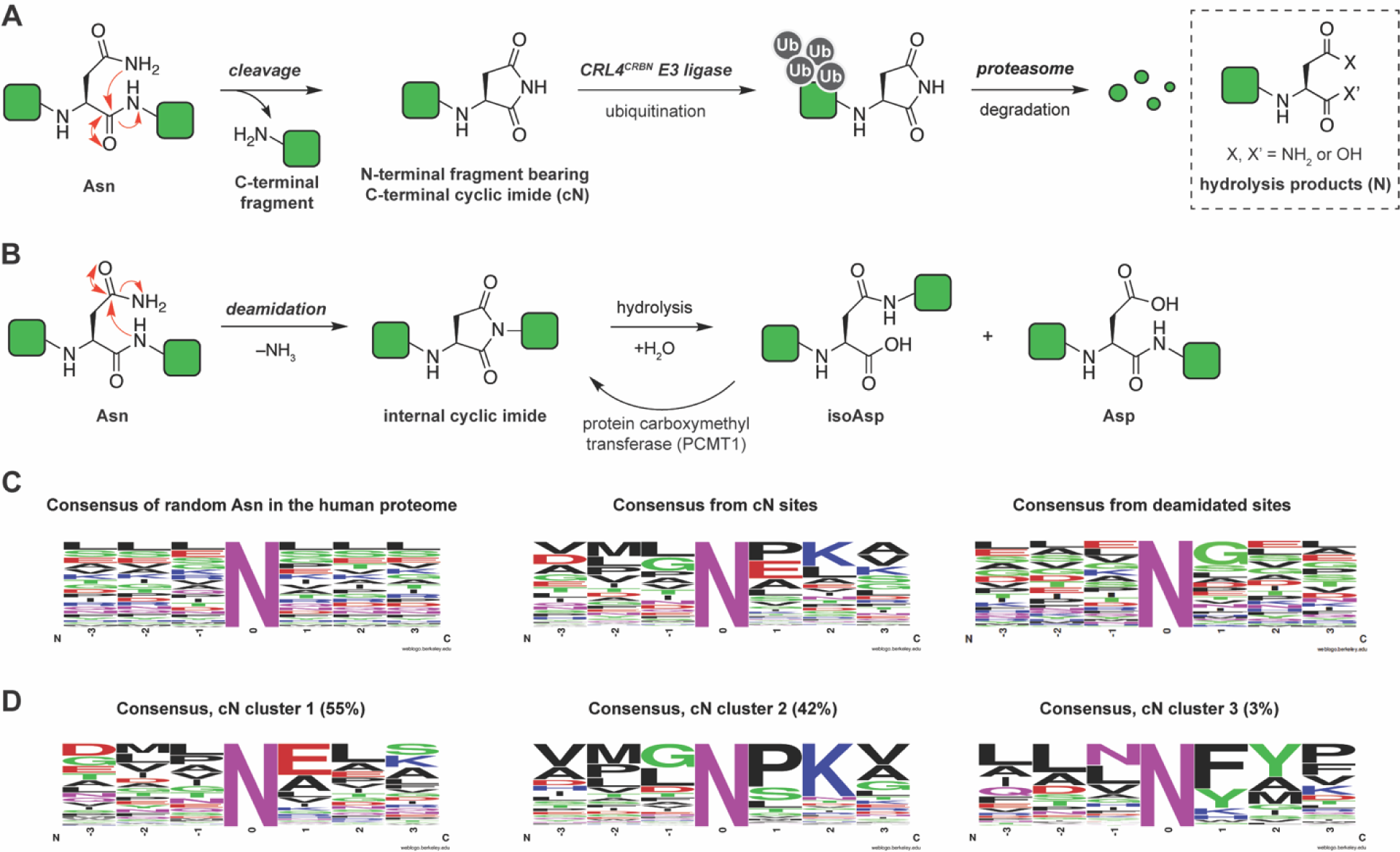
Regulatory mechanisms and consensus sequences of asparagine residues that are susceptible to cleavage or deamidation. **(A)** Schematic of the formation and regulation of C-terminal cyclic imide modifications. **(B)** Schematic of the regulation of protein damage arising from deamidation. **(C)** Frequency charts of the sequence alignment spanning the –3 to +3 residues flanking random asparagine sites extracted from the human proteome, C-terminal aspartimide sites or deamidation sites observed in CPTAC datasets. **(D)** Frequency charts of the sequence alignment for the three clusters generated from hierarchical clustering of the +1 and +2 residues of the cN sites observed in CPTAC datasets based on similarity of amino acid properties. The percentage of each cluster out of the total population is noted. All alignments were generated with weblogo.berkeley.edu.

The cell maintains quality control mechanisms to repair or remove damaged proteins(21). One of the best elucidated repair mechanisms is the conversion of isoaspartyl residues, the end product of asparagine deamidation, to L-aspartate by protein O-carboxymethyltransferase (PCMT1), a conserved enzyme that partially repairs isoaspartate (**Fig. 1B**). The loss of PCMT1 is associated with accelerated aging(22) and disorders such as growth retardation and epileptic seizures that may be due to the dramatic increase in isoaspartate-containing proteins(23, 24), suggesting that the aberrant buildup of damaged proteins can cause highly deleterious effects. Recently, we discovered that protein damage from C-terminal cyclic imide modifications is endogenously targeted by the E3 ligase substrate adapter cereblon (CRBN)(25). Protein substrates bearing the C-terminal cyclic imide are recognized by CRBN and are subsequently degraded via the ubiquitin-proteasome system (**Fig. 1A**). Thalidomide and its derivatives, widely used therapeutic agents, mimic the C-terminal cyclic imide modification and therefore may perturb the turnover of these endogenous substrates(25, 26). As mutations of CRBN have been associated with intellectual disability(27, 28) and other neurological disorders reminiscent of those connected to the loss of PCMT1, the biological effects following accumulation of protein damage from C-terminal cyclic imides and deamidation may be similar. However, while there is a strong understanding of the intrinsic molecular features of the primary and secondary structure of proteins that give rise to deamidation(29–32), the factors that affect the formation of C-terminal cyclic imides have not been fully elucidated. Here, we characterize the formation of C-terminal cyclic imides via intramolecular cleavage of peptides and proteins in vitro and in cells, their recognition by CRBN, and the functional consequence of this form of protein damage on the proteome.

## Results

### The protein damage C-terminal cyclic imide consensus sequence

Previously, we mapped thousands of semi-tryptic cN and cQ modification sites by meta-analysis of Clinical Proteomic Tumor Analysis Consortium (CPTAC) datasets from primary human tissues and the NCI7 cell line panel(25). Although these modifications could emerge in situ during mass spectrometry analysis(16), we observed consensus sequences that are predicted to be more sensitive to C-terminal cyclic imide formation. In contrast to a background sequence alignment generated by a random selection of 50,000 asparagine sites from the human proteome, the alignment of sequences flanking mapped C-terminal aspartimide sites revealed an enrichment of certain residues at the +1 and +2 positions of cN (**Fig. 1C, Dataset S1**). Notably, P appears as the most frequent residue at the +1 position for cN, likely because it hinders the deamidation process where the backbone amide attacks the side chain carbonyl (**Fig. 1B**) and in turn favors the attack of the backbone carbonyl by the side chain amide (**Fig. 1A**). Charged residues such as E and K, which may facilitate the proton transfer or nucleophilic addition, are also enriched at the +1 and +2 positions of the consensus sequence, respectively. Meta-analysis of several of the same public proteomics datasets (NCI7, brain, and breast)(33–35) for asparagine deamidation using a similar workflow revealed an enrichment of G at the +1 position (**Fig. 1C, Dataset S2**), mirroring the reported sequence preference for faster deamidation kinetics(36, 37) and demonstrating the difference in consensus sequence between C-terminal aspartimide cleavage sites and deamidation sites. A similar comparison of the consensus sequence of mapped cQ sites to randomly selected glutamine residues revealed minor but less notable shifts in the consensus sequence relative to randomly selected glutamine residues (**Fig. S1A**).

Hierarchical clustering of the +1 and +2 residues of the identified cN sites based on the similarity of amino acid properties, which yielded three clusters. In the NEX cluster, accounting for more than half of the total population, E is the most enriched at the +1 position and is followed by neutral residues at the +2 position (**Fig. 1D**). In contrast, the second major cluster NXK had a significant enrichment of K at the +2 position and P at the +1 position (**Fig. 1D**). We thus experimentally investigated the two major consensus sequences of cN derived from this meta-analysis.

## Peptide and protein primary sequence influences C-terminal cyclic imide formation

We synthesized a library of peptides that contain asparagine in the middle and varying amino acid residues at the +1 or +2 position to examine their propensity for cN formation. These peptides possess a common N-terminal VEALLNXXALE or VEALLNXXALLE consensus sequence, where X is the variable +1 or +2 positions. This sequence was designed so that each position is represented by one of the most frequent amino acids in the predicted consensus and the resulting peptides are synthetically tractable and detectable by mass spectrometry. The peptides were incubated under physiological conditions for 2 days to generate the C-terminal aspartimide fragment (VEALLcN). The quantification of VEALLcN and its hydrolysis products was performed by dividing the peak area of the lightest isotope of the modified species by that of the parent peptide to exclude the contribution from deamidation. Investigation of peptides that model the NEX consensus sequence revealed that neutral residues at the +2 position promoted the formation of cN and its hydrolysis products (N) more than positively charged residues K and R (**Fig. 2A, Fig. S1B, Dataset S3**).

**Figure 2.**
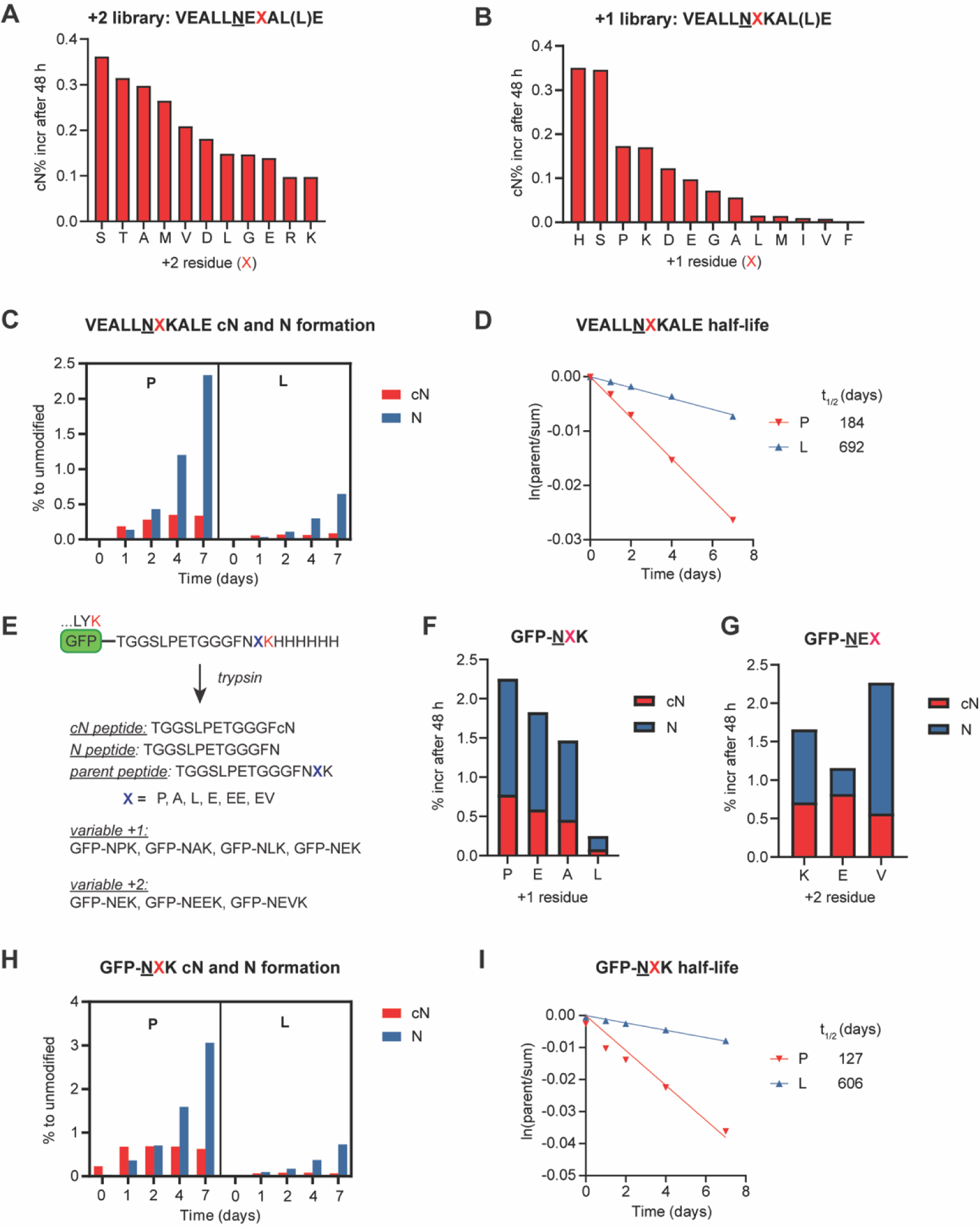
Impact of primary sequences of peptides and proteins on C-terminal cyclic imide formation. **(A–B)** Quantification of cN formation with a peptide library carrying variable +2 **(A)** or +1 **(B)** residues after 48 h incubation in 100 mM Na_2_HPO_4_ (pH 7.4) at 37 °C. **(C)** Time-course formation of cN and N from the indicated peptides in 100 mM Na_2_HPO_4_ (pH 7.4) at 37 °C. **(D)** Derivation of the half-life for the indicated peptides in 100 mM Na_2_HPO_4_ (pH 7.4) at 37 °C. **(E)** Design of engineered GFPs to evaluate the impact of protein primary structure on intramolecular cleavage. **(F–G)** Quantification of cN and N formation from GFP model proteins with varying +1 **(F)** or +2 **(G)** residues after 48 h incubation in 100 mM Na_2_HPO_4_ (pH 7.4) at 37 °C. **(H)** Time-course formation of cN and N from the indicated proteins in 100 mM Na_2_HPO_4_ (pH 7.4) at 37 °C. **(I)** Derivation of the half-life for the indicated proteins in 100 mM Na_2_HPO_4_ (pH 7.4) at 37 °C.

To examine the NXK consensus, we measured peptides with K at the +2 position and a variable +1 residue for formation of the VEALLcN fragment. Incubation of these peptides for 48 h at 37 °C, pH 7.4 revealed that small, polar residues such as H, S, and P facilitated cN and N formation, while bulky, hydrophobic residues such as V and F greatly impeded cleavage, with an approximately 200-fold difference in cN formation rate across the library (**Fig. 2B, Fig. S1C**). Moreover, for both variable +1 and +2 libraries, the increase in the hydrolysis products highly correlated to that of cN formation (**Fig. S1D–E**), suggesting that cleavage at asparagine transitions through the C-terminal cyclic imide modification. Importantly, longer incubation of two representative peptides bearing P or L at the +1 position revealed that the steady accumulation of the hydrolysis product and corresponding decrease in the parent peptide over time are faster for the peptide with P at the +1 position (**Fig. 2C**). The faster kinetics in cN formation in turn led to a shorter half-life of the parent peptide (**Fig. 2D**). These observations experimentally validate the two distinct consensus sequences predicted by meta-analysis of the proteomics datasets.

Separately, we examined the variation of residues at the –1 position relative to two representative peptides with the consensus sequence VEALXNESALE or VEALXNALALE. Although analysis may be complicated by differences in the detection efficiency of the VEALXcN species by mass spectrometry, these data showed that L and P promoted cN formation in the context of the first peptide but were less impactful in the context of the second peptide (**Fig. S1F**).

We next evaluated if the determinants of peptide cleavage propensity extended to proteins by generating GFP proteins modified at the C-terminus to display the most sensitive sequences observed from the peptides (**Fig. 2E**). Tryptic digestion sites were also incorporated to yield peptide fragments with facile ionization and detection on the mass spectrometer. After incubation under physiological conditions for 48 h, the engineered GFPs were digested and the resulting parent peptide and semi-tryptic cN and N peptides representing the cleavage products were detected using selected ion monitoring (SIM) for quantification. As before, the +1 position influenced the rate of protein cleavage at asparagine, whereas variation of the +2 position had a moderate impact on the protein cleavage kinetics (**Fig. 2F–G**), verifying the translatability of the consensus sequence to unstructured region of proteins. Moreover, a time course incubation of GFP-NPK and GFP-NLK proteins confirmed that C-terminal cyclic imide formation correlates with the buildup of hydrolysis products and reduction in the parent peptide, resulting in a shorter half-life of the intact protein (**Fig. 2H–I**). Although mass spectrometry analyses can generate C-terminal cyclic imide species in situ(16, 17), the consensus sequences for these modifications derived from proteomic datasets were largely validated in these controlled experimental measurements on defined peptides and proteins. Thus, these data collectively establish the primary sequences that are more sensitive to C-terminal cyclic imide formation, which were predicted by global proteomics datasets and are distinct from those sensitive to deamidation.

### Asparagine is more susceptible to cleavage than glutamine

To compare the cleavage kinetics between asparagine and glutamine residues, we substituted asparagine in the most susceptible sequences identified from the peptide library with glutamine and incubated them under the same physiological conditions. Glutamine-containing peptides possessed slower kinetics in the C-terminal cyclic imide formation and therefore were significantly more stable to intramolecular cleavage (**Fig. 3A, Dataset S4**), resulting in their commensurately longer half-lives than the asparagine-containing peptides (**Fig. S2A–B**). These data align with the favorable kinetics of five-membered over six-membered ring formation(38). Similarly, asparaginyl peptides are more sensitive to deamidation and have more than 100-fold shorter half-life than glutaminyl peptides(37). Thus, we primarily focused on the characterization of intramolecular cleavage at asparagine in our subsequent analyses.

**Figure 3.**
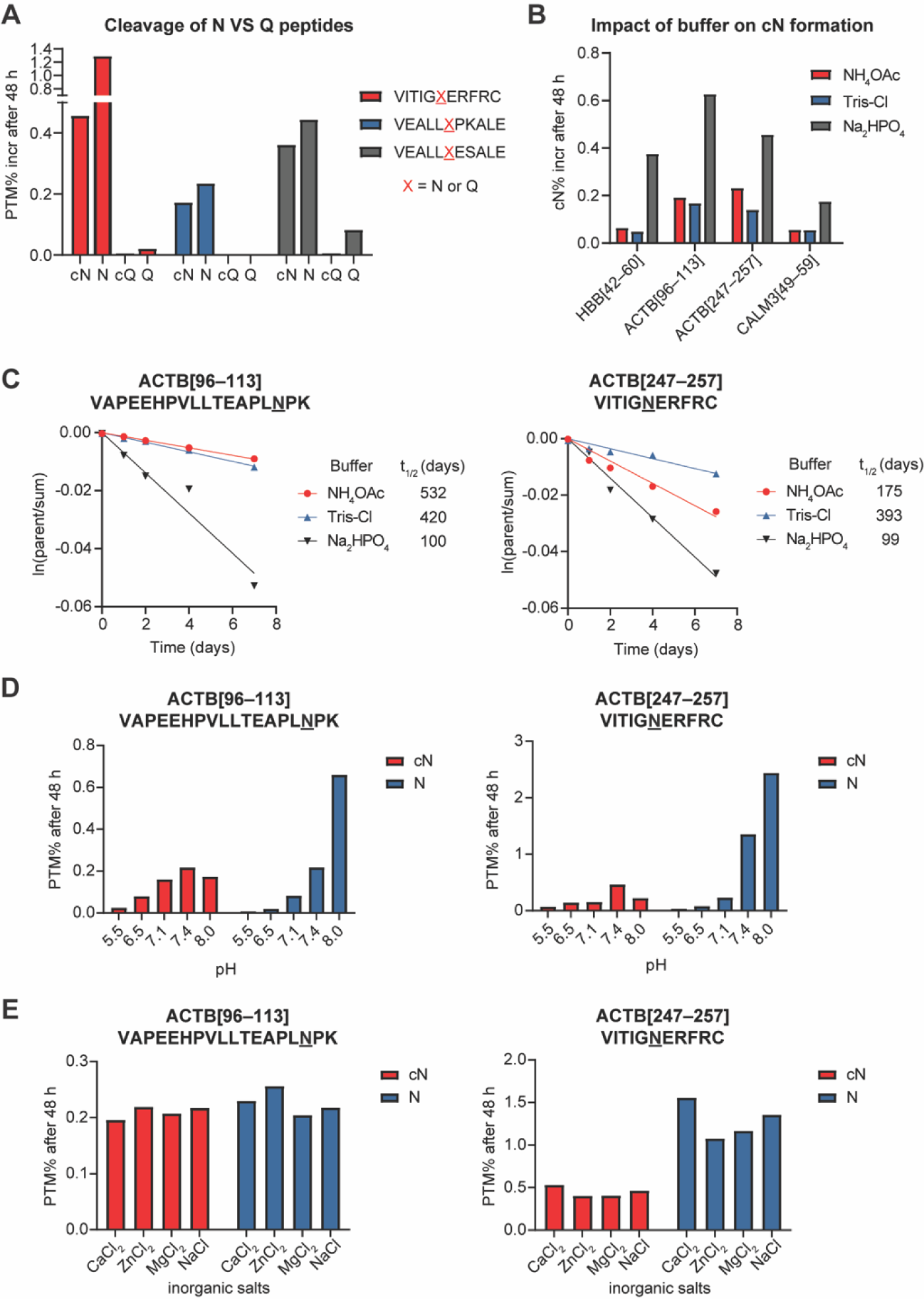
Effect of buffer properties on C-terminal cyclic imide formation. **(A)** Comparison of C-terminal cyclic imide and hydrolysis products for peptides with a differing central N or Q residue after 48 h incubation in 100 mM Na_2_HPO_4_ (pH 7.4) at 37 °C. **(B)** Quantification of cN and N formation from four representative peptides after 48 h incubation in 100 mM NH_4_OAc, Tris-Cl, or Na_2_HPO_4_ (all pH 7.4) at 37 °C. **(C)** Time-course and half-life derivation for the indicated peptides in 100 mM NH_4_OAc, Tris-Cl or Na_2_HPO_4_ (all pH 7.4) at 37 °C. **(D–E)** Quantification of cN and N species from the indicated peptides after 48 h incubation in 100 mM Na_2_HPO_4_ with variable pH at 37 °C with variable pH **(D)** or containing 1 mM inorganic salts at pH 7.4 **(E)**.

### Influence of buffer on intramolecular cleavage

We next examined the external factors that affect the rate of C-terminal cyclic imide formation and peptide cleavage in vitro. Since deamidation kinetics are notably accelerated 2- to 3-fold in the presence of phosphates and other buffers that influence the rate of cyclization(39, 40), we examined whether buffer properties would similarly promote the rate of peptide cleavage. Four peptides derived from genes HBB, ACTB and CALM3 carrying sequences of the most frequent cN sites from the global proteomics meta-analysis were incubated in buffers with ammonium, Tris, or phosphate ions. Based on isotopic patterns, negligible deamidation was observed on these peptides, indicative of distinct sequence-dependent forms of protein damage at Asn. For all peptides tested, phosphate buffer elevated cN formation by 2- to 7-fold (**Fig. 3B**) and gave rise to a correspondingly shorter half-life of the parent peptide was measured (**Fig. 3C, Fig. S2C**). The hydrogen bonding between the phosphate group and the electrophilic carbonyl of the backbone may therefore promote proton transfer and accelerate the formation of the reaction intermediate for C-terminal cyclic imide formation, an effect similar to that present in internal cyclic imide formation during deamidation.

The nucleophilic addition and excision of the leaving group involved in the C-terminal cyclic imide formation are also subject to acid-base catalysis. We therefore tested different pH values within the working range of sodium phosphate buffer and observed the least reactivity at acidic pH, the most cN formation at pH 7.4 and the most hydrolysis product formation at pH 8.0 for both evaluated peptides (**Fig. 3D**). This demonstrates high sensitivity to pH for both the formation and hydrolysis of the C-terminal cyclic imide. Additionally, we explored the effect of metal ions that may serve as Lewis acids or redox centers and catalyze many chemical transformations in biological systems(41). We evaluated the most common metal ions that participate in biological catalysis, zinc, magnesium, and iron, but none of these metal ions significantly affected the rate of C-terminal cyclic imide formation or hydrolysis (**Fig. 3E**). These observations reveal that the degradation of asparagine-containing primary sequences via intramolecular cleavage is promoted by phosphate ions and basic conditions but is unaffected by metal ions.

### Secondary protein structural features impact cleavage

In addition to the primary sequence, the secondary structure of the protein can influence the cleavage event at a given asparagine residue(42). As the consensus sequence derived from the meta-analysis is broadly predictive of sensitivity to deamidation or intramolecular cleavage, we next computationally analyzed the most frequently modified residues in the context of the protein secondary structure to shed light on the protein structural features characteristic of asparagine sites susceptible to both forms of protein damage. We first generated two background datasets by random selection of 5000 asparagine sites from the SwissProt human proteome for comparison to the 2000 most frequent sites for intramolecular cleavage or deamidation identified from meta-analysis by peptide spectral match (PSM) counting. For the datasets “random 1”, “random 2”, “deamidation” and “cleavage”, we extracted structural information by ChimeraX(43) from either experimentally determined structures that contain the site and have resolution ≤ 3.5Å, or high-confidence structures predicted by AlphaFold(44) (site pLDDT ≥ 70) if experimental structures were unavailable (**Fig. 4A, Dataset S5**). The compilation of relative solvent-excluded surface area (relSESA) of the asparagine sites, which is defined as the surface area of the asparagine in the structure normalized to a reference surface area of the asparagine in a Gly-Asn-Gly tripeptide, revealed a strikingly lower surface exposure of cleavage sites when compared to the two random datasets, which had no statistically significant difference from each other, and the deamidation dataset, which had a slightly higher surface exposure (**Fig. 4B–C**). Moreover, Asn residues prone to cleavage have the shortest distance between the side chain amide and the backbone carbonyl out of the four groups (**Fig. 4D–E**), which likely promotes the initial nucleophilic addition of the side chain. We additionally computed the backbone dihedral angles psi and phi of the asparagine sites and visualized the distribution of conformations on Ramachandran plots(45). The distributions of the two background asparagine datasets were similar to each other (**Fig. S3A**). By contrast, the distribution of cleavage sites showed slight enrichment of right-handed alpha helix (3^rd^ quadrant) and depletion of left-handed alpha helix (1^st^ quadrant) in comparison to deamidation sites (**Fig. S3B**), highlighting the distinct regions of the proteome that are more sensitive to intramolecular cleavage or deamidation as compared to the average asparagine residue.

**Figure 4.**
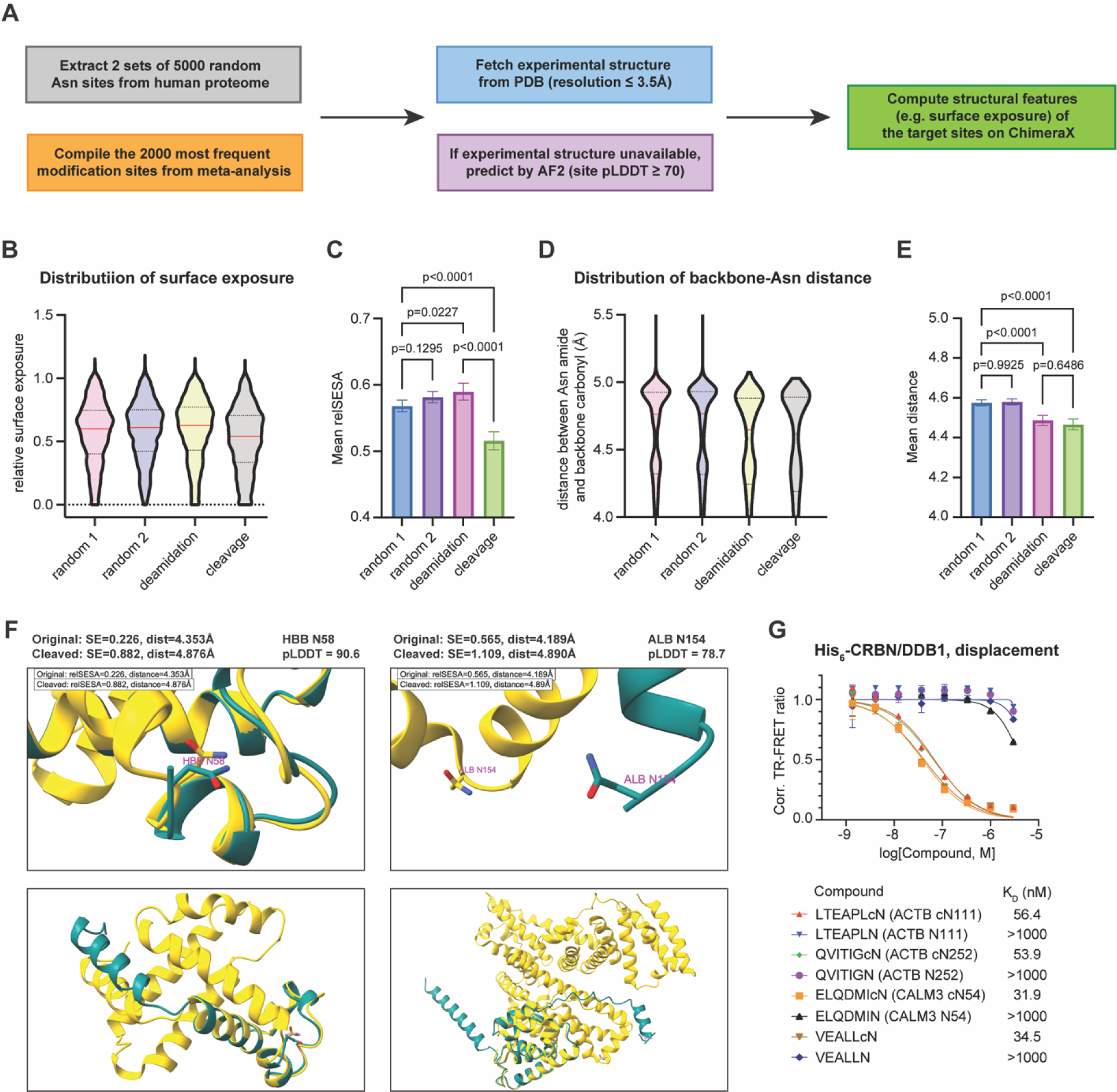
Influence of protein secondary structures on the propensity for cleavage. **(A)** Workflow of the computational analysis to derive protein secondary structure features. **(B)** Distribution of relative surface exposure for frequently observed deamidation and cleavage sites in comparison to two background datasets comprised of randomly extracted Asn sites. Median values are indicated by a red line and the 25% and 75% percentiles by dotted lines. **(C)** Comparison of mean values of relative surface exposure across the four evaluated groups. **(D)** Distribution of distance between Asn amide and backbone carbonyl for frequent deamidation and cleavage sites in comparison to two background datasets comprised of randomly extracted Asn sites. Median values are indicated by a red line and the 25% and 75% percentiles by dotted lines. **(E)** Comparison of mean values of distance across the four evaluated groups. All comparisons were performed using a one-way ANOVA with Šidák’s multiple comparisons test and p-values are noted. **(F)** Overlay of the original experimental structure in yellow and the predicted structure after cleavage at the indicated asparagine residue in teal for HBB N58 and ALB N154. The calculated values of surface exposure and distance and the pLDDT scores for the prediction of asparagine cleavage sites are noted. **(G)** Dose titration of the indicated peptides in TR-FRET assay against the His_6_-CRBN/DDB1 complex with the determined K_D_ values noted. The experiment was performed with 3 technical replicates.

We extended the analysis of asparagine conformations to intein splicing, one of the most studied mechanisms of intramolecular cleavage at asparagine(46). We derived the same parameters from the structure of the branched intermediate of the Mxe GyrA intein(47), where the asparagine side chain amide is primed for the nucleophilic attack on the backbone carbonyl at the splice junction, as well as the structure of the Mxe GyrA intein after splicing(48). The key asparagine residue in the intein had a lower relSESA and was closer to the backbone carbonyl in the branched intermediate as compared to the C-terminal aspartimide after intein excision (**Fig. S3C**), although the relative inaccessibility of the C-terminal aspartimide on the intein may still block recognition by CRBN(25). To evaluate if C-terminal aspartimides generated from protein damage events in the human proteome are more solvent exposed, we selected nine of the most frequently occurring truncation fragments to model by AlphaFold and extracted structural information of the six high confidence predictions (pLDDT ≥ 70) by ChimeraX for comparison to the full-length protein (e.g., HBB[1–58N] from truncation of full-length HBB[1–147]). Five out of the six evaluated fragments showed a spike in both relSESA and distance between the side chain amide and adjacent carbonyl in comparison to their parent structures (**Fig. 4F, Fig. S3D**). Collectively, these computational analyses shows that Asn residues in buried or strained regions of proteins can promote the nucleophilic attack required for protein cleavage, but after cleavage the resulting C-terminal aspartimide is solvent exposed to facilitate recognition by CRBN.

To assess recognition of the C-terminal cyclic imide peptides by CRBN, we synthesized three heptamer peptides representing sequences surrounding the most frequent cN sites with cyclic or uncyclized asparagine and measured their binding by competitive displacement time-resolved Förster resonance energy transfer (TR-FRET) assay using the His_6_-CRBN/DDB1 complex labeled with CoraFluor-1 as the FRET donor(49, 50) (**Fig. S3E**). We assessed the saturation binding of tracer molecule thalidomide-fluorescein isothiocyanate (thal-FITC) to the His_6_-CRBN/DDB1 complex to derive the tracer equilibrium dissociation constant (K_D_) to convert ligand IC_50_ to K_D_ values (**Fig. S5F–G**). The evaluation of the three cN peptides and peptide library standard (VEALLcN) showed that CRBN has a rather flexible substrate scope for cN, but does not bind to the identical sequences bearing uncyclized asparagine (**Fig. 4G**). These data indicate that CRBN recognizes sequences that endogenously form the C-terminal cyclic imide upon cleavage.

### Influence of external stressors on protein cleavage

The cell encounters a variety of environmental and metabolic stressors, such as reactive oxygen species (ROS), heat stress, mechanical stress and dysregulated pH levels, which have been connected to accelerated protein damage and compromised proteostasis due to misfolding and aggregation(51–54). To assess the effect of stressors on cleavage, we subjected the GFP-NPK model protein to high temperature, hydrogen peroxide or vortexing to simulate heat, oxidative or mechanical stress, respectively, and observed a dramatic increase in the cleavage products upon heat stress (**Fig. 5A, Dataset S6**). Formation of the cN fragment and its hydrolysis products was further accelerated by higher pH and temperature (**Fig. 5B–C**). Notably, the hydrolysis products particularly increased under physiologically relevant conditions, including elevation of pH from 6.5 to 7.1 and temperature from 22 °C to 37 °C.

**Figure 5.**
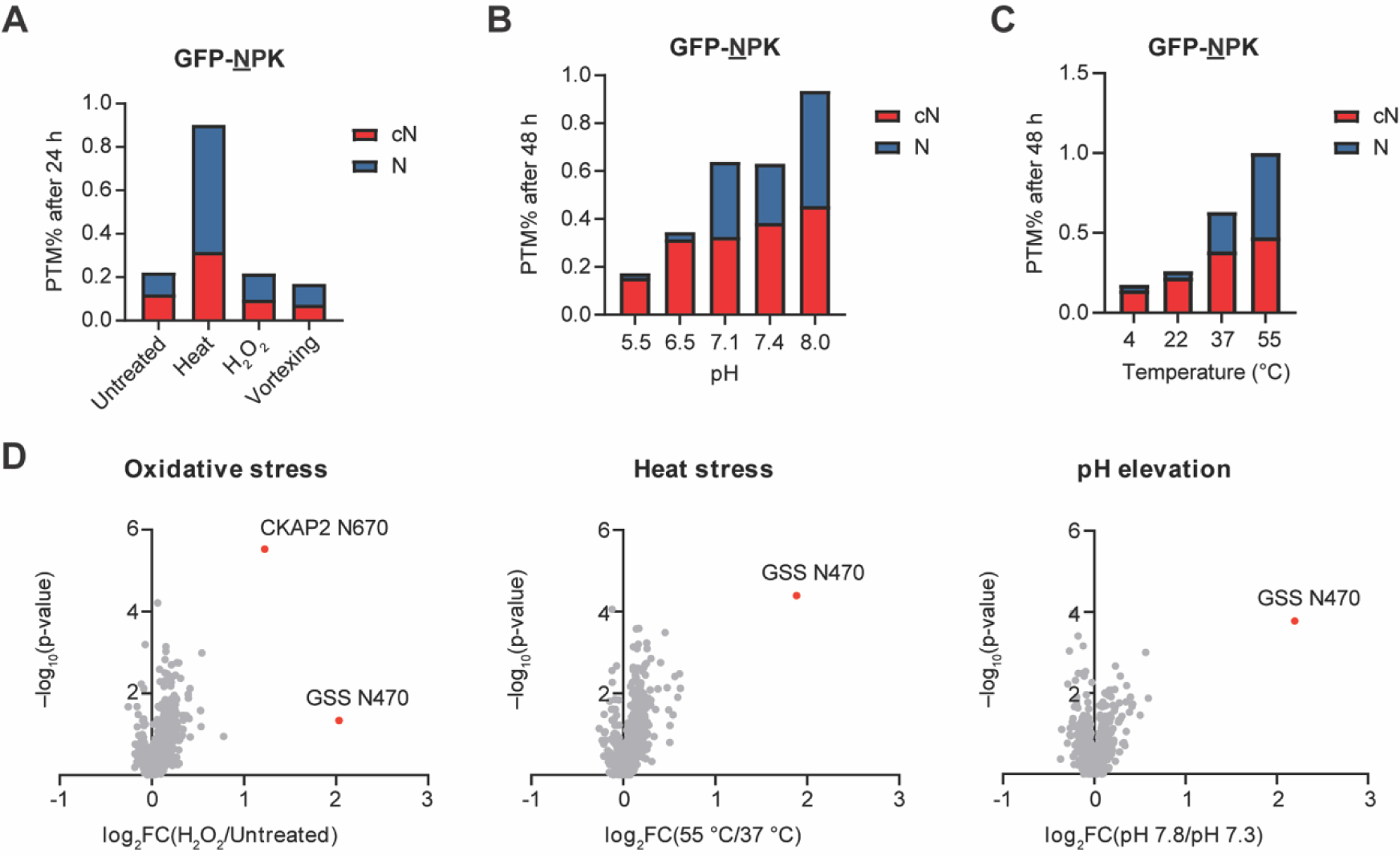
Effect of external stressors on cleavage in vitro and in cells. **(A)** Quantification of cN and N species from the GFP model protein after 24 h exposure to heat stress (55 °C), 100 µM H_2_O_2_, or vortexing. The protein was incubated in 100 mM Na_2_HPO_4_ buffer, pH 7.4 at 37 °C unless otherwise noted. **(B–C)** Quantification of cN and N species from the GFP model protein after 48 h incubation in 100 mM Na_2_HPO_4_ buffer and variable pH at 37 °C **(B)** or at pH 7.4 with variable temperature **(C)**. **(D)** Volcano plots of semi-tryptic peptides ending with N in CRBN-KO HEK293T treated with 100 µM H_2_O_2_ for 40 min, incubated at 55 °C for 40 min, or incubated at pH 7.8 for 4 h compared to untreated cells incubated at pH 7.3, 37 °C. The experiment was performed with 4 biological replicates. P-values for the abundance ratios were calculated by one-way ANOVA with TukeyHSD post-hoc test.

To globally map changes in cleavage events upon exposure to common stressors, we challenged HEK293T cells with CRBN knockout (KO) by CRISPR/Cas9 to brief oxidative stress, heat stress, or pH elevation. Employing tandem mass tag (TMT)-based quantitative proteomics, we quantified 629 semi-tryptic peptides ending with N, which are indicative of intramolecular cleavage events and found more than half of these peptides were elevated under all stress conditions (**Fig. 5D**). Despite the slow kinetics of C-terminal glutarimide formation in vitro, we measured 189 semi-tryptic peptides ending with Q although overall changes in abundance were less apparent compared to those ending with N (**Fig. S4A**). Additionally, 5455 peptides bearing at least one deamidated asparagine residue were quantified, more than half of which were similarly upregulated under all stress conditions (**Fig. S4B**). The sequence alignments for the deamidation and C-terminal cyclic imide sites identified from this proteomics experiment showed enrichment of residues similar to those derived from the meta-analysis, indicating reproducibility of the consensus sequences (**Fig. S4C**). Collectively, these data demonstrate that heat, pH, and oxidative stress accelerate protein damage and formation of the C-terminal cyclic imide in vitro and in cells.

### GSS is regulated by CRBN upon cleavage

In the stress proteomics data, we identified the semi-tryptic peptide from glutathione synthetase (GSS) representing a cleavage event close to the C-terminus at N470 as the most significantly upregulated cleavage site across the tested stressor conditions, while the total GSS protein level was unaltered (**Fig. 5D, Fig. S4D**). The active form of GSS is a homodimer with subunits composed of 474 residues and is responsible for catalyzing the last step of the biosynthesis of glutathione (GSH), a prevalent antioxidant that protects cells from oxidative damage and participates in many biological pathways. Patients with hereditary GSS deficiency exhibit reduced GSH levels and, in severe cases, dysfunction of the central nervous system(55). Genetic studies of individuals with GSS deficiency have mapped different missense mutations in the coding sequence that are proximal to the identified N470 cleavage site, including G464V and D469E(56), indicating the structural and functional importance of the C-terminal region of GSS (**Fig. S5A**). GSS N470 is followed by P at the +1 position which is predictive of cleavage based on our primary sequence analysis. Furthermore, GSS N470 and the rest of the C-terminal region are buried in the crystal structure(57), but N470 is predicted to become more solvent-exposed after cleavage (**Fig. S5B**) and hence more readily recognized by CRBN. Thus, if GSS is subject to intramolecular cleavage at N470 during aging or upon stress, this may render the protein dysfunctional and thus require recognition by CRBN for ubiquitination and proteasomal degradation of the damaged and inactivated protein (**Fig. 6A**).

**Figure 6.**
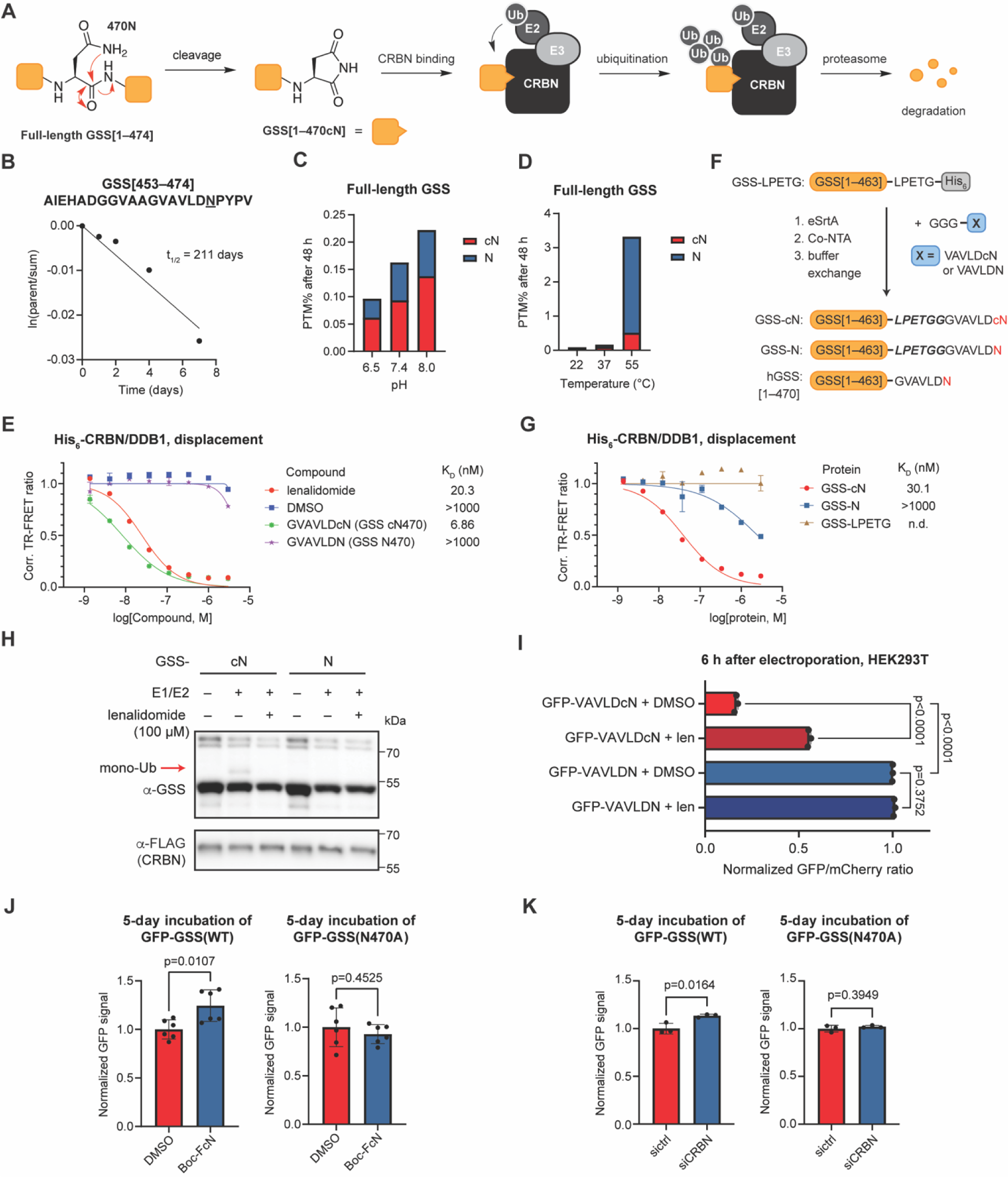
CRBN regulates GSS upon cleavage. **(A)** Schematic of the cleavage at N470, recognition by CRBN and removal by the ubiquitin-proteasome system of GSS. **(B)** Derivation of the cleavage half-life for the GSS parent peptide in 100 mM Na_2_HPO_4_ (pH 7.4) at 37 °C. **(C)** Quantification of cN and N species from full-length GSS protein after 48 h incubation in 100 mM Na_2_HPO_4_ buffer at the indicated pH at 37 °C. **(D)** Quantification of cN and N species from full-length GSS protein after 48 h incubation in 100 mM Na_2_HPO_4_ (pH 7.4) at the indicated temperature. **(E)** Dose titration of the indicated compounds in TR-FRET assay against the His_6_-CRBN/DDB1 complex with the determined K_D_ values noted. **(F)** Schematic of sortase system used to generate tagged GSS from GSS-LPETG-His_6_ and the comparison to the native sequence of human GSS (hGSS). **(G)** Dose titration of GSS protein tagged with C-terminal cyclic or acyclic asparagine in TR-FRET assay against the His_6_-CRBN/DDB1 complex with the determined K_D_ values noted. All TR-FRET experiments were performed with 3 technical replicates. **(H)** In vitro ubiquitination of GSS protein tagged with C-terminal cyclic or acyclic asparagine with K0 ubiquitin. **(I)** Flow cytometry analysis of the GFP levels in HEK293T cells 6 h after electroporation of GFP tagged with the GSS sequence bearing cyclic or acyclic C-terminal asparagine. The experiment was performed with 3 biological replicates. Comparisons were performed using a one-way ANOVA with Šidák’s multiple comparisons test and p-values are noted. **(J)** Quantification of GFP level in cells stably expressing GFP fusion of full-length wild-type GSS or GSS bearing N470A mutation by flow cytometry. Cells were treated with DMSO or 200 µM Boc-FcN across 6 biological replicates for 5 days with media renewal every 2 days. **(K)** Quantification of GFP level in cells stably expressing GFP fusion of full-length wild-type GSS or GSS bearing N470A mutation by flow cytometry. Cells were transfected with 30 nM non-targeting siRNA or siRNA targeting CRBN with 3 biological replicates and incubated for 5 days post-transfection. Comparisons for the reporter cell lines were performed using an unpaired two-tailed t-test and p-values are noted.

Indeed, incubation of the GSS C-terminal parent peptide (GSS[453–474]) and the full-length GSS protein revealed that cleavage of the peptide at N470 has an intrinsic half-life of 211 days, which is more stable in the context of the protein under normal physiological conditions (**Fig. 6B, Fig. S5C, Dataset S7**). However, exposure of the GSS protein to high pH or temperature dramatically increased the rate of cleavage and generation of cN and N at this site (**Fig. 6C–D**), which validates the sensitivity of this site to stress in the cellular context.

To confirm that CRBN recognizes the C-terminal aspartimide on GSS in the peptide and protein context, we synthesized the GSS[464–cN470] heptamer peptide and evaluated its binding to CRBN by TR-FRET displacement assay. We measured K_D_,_GSS[464–cN470]_ to be 6.86 nM, which is 3-fold better than lenalidomide (**Fig. 6E**). In contrast, the identical sequence bearing acyclic asparagine had no measurable affinity towards CRBN. Our subsequent efforts to assess the functional engagement by CRBN of GSS[1–cN470] were hindered by aggregation of the product during the traceless generation of GSS[1–cN470] using protein ligation methods. We thus used the sortase system to semi-synthetically install the cyclized or uncyclized asparagine to GSS-LPETG-His_6_, affording the products GSS-cN or GSS-N consisting of with a 6-AA (LPETGG) scar on the otherwise intact native human GSS sequence (**Fig. 6F**). The cN- and N-tagged GSS proteins were first evaluated for CRBN binding, where GSS-cN showed comparable K_D_ to the peptide ligand while GSS-N did not bind to CRBN (**Fig. 6G**). A complementary TR-FRET assay measuring the direct binding of the sortase-reacted GFP (**Fig. S5D**) and CRBN showed a much higher affinity for the GFP tagged with the GSS cN peptide (GFP-VAVLDcN) compared to the acyclic GFP-VAVLDN control and GFP tagged with FcN or FcQ, validating the tight binding between the GSS sequence and CRBN (**Fig. S5E–F**). The GSS proteins were further subjected to an in vitro ubiquitination assay using K0 ubiquitin in the presence of the CRL4^CRBN^ complex isolated from HEK293FT cells stably expressing FLAG-CRBN. The specific ubiquitination band was only observed for GSS-cN and diminished by addition of lenalidomide (**Fig. 6H, Fig. S6A**). This specific ubiquitination event was similarly observed for GFP-VAVLDcN (**Fig. S6B**). Moreover, we observed the rapid degradation of GFP-VAVLDcN in contrast to GFP-VAVLDN 6 h after electroporation into HEK293T cells, which was rescued by co-treatment with lenalidomide (**Fig. 6I**), verifying the functional engagement of the GSS sequence by CRBN in vitro and in cells.

To sensitively monitor if GSS levels change in response to this intrinsic cleavage event in a cellular context, we constructed reporter cell lines in HEK293T that stably express either GFP-GSS(WT) or GFP-GSS(N470A) containing alanine mutation at the cleavage site (**Fig. S6C**). The reporter cell lines were treated with Boc-FcN or siRNA targeting CRBN over 5 days to temporally inhibit recognition by CRBN, after which the level of GFP-GSS was measured by flow cytometry. We expected that the inhibition of CRBN would reduce degradation of the GSS cleavage products formed at N470 for GFP-GSS(WT) cells while the level of GFP-GSS(N470A), which cannot form the cyclic imide degron at this position, would be unaffected. Indeed, we observed a 24% increase in GFP signal on average after Boc-FcN treatment relative to DMSO-treated controls in cells expressing GFP-GSS(WT) while there was no significant change between treatment groups in cells expressing GFP-GSS(N470A) (**Fig. 6J**). Likewise, upon CRBN knockdown for 5 days (**Fig. S6D**), a 14% increase on average from controls was observed for GFP-GSS(WT) reporter cells but not GFP-GSS(N470A) cells (**Fig. 6K**), congruent with the model where the C-terminal cyclic imides generated via cleavage are removed by CRBN in a timely manner and thus their accumulation is commuted. Collectively, these data demonstrate that the C-terminal cyclic imide modification formed at the susceptible cleavage site of GSS is recognized and regulated by CRBN.

### Protein aggregation following intramolecular cleavage

As other forms of protein damage that are known to elicit deleterious effects on the modified proteins(58, 59), C-terminal cyclic imides generated from intramolecular cleavage may disrupt the protein homeostasis and trigger aggregation, which have been linked to severe biological consequences such as neurodegenerative diseases(60). To examine the effect of intramolecular cleavage on the stability of GSS, we recombinantly expressed GSS[1–470] (cleaved) and GSS[1– 474] (full-length) with an N-terminal His_6_ tag under the same conditions and purified them using affinity chromatography. The analysis of two constructs at the same concentration by protein size-exclusion chromatography (SEC) revealed an increased population exhibiting higher molecular weight in GSS[1–470] compared to GSS[1–474] (**Fig. 7A**), which may stem from the aggregation of the cleaved protein. The SDS-PAGE analysis of the two constructs confirmed the presence of higher molecular weight bands in GSS[1–470] (**Fig. S7A**). Apparent white precipitation was also observed for GSS[1–470] but not GSS[1–474] after three freeze-thaw cycles of the aliquots at the same concentration, indicating the truncated protein is unstable (**Fig. S7B**). We additionally evaluated the GSS constructs by PROTEOSTAT staining(61) which enables the quantitative analysis of the amount of aggregation by fluorescence reads. At all evaluated concentrations, GSS[1–470] exhibited more fluorescence than GSS[1–474] (**Fig. 7B, Dataset S8**), indicative of a higher propensity for aggregation. Collectively, these data demonstrate that the intramolecular cleavage of GSS at N470 facilitates its oligomerization and aggregation.

**Figure 7.**
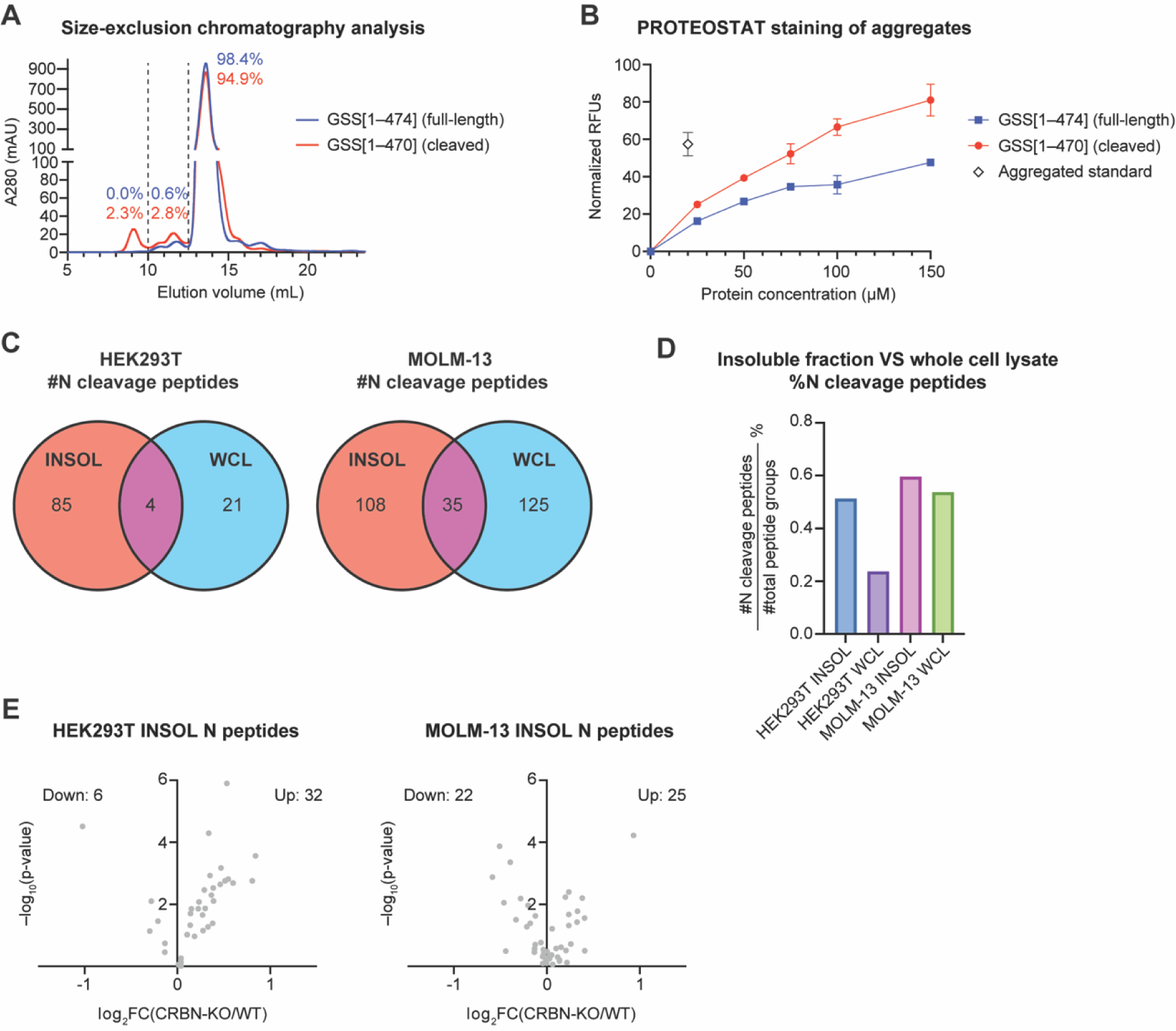
Association between cleavage products and protein aggregation. **(A)** Size-exclusion chromatography of the affinity-purified GSS proteins. Percentages of the peak areas are noted for different ranges of elution volumes. **(B)** Quantification of aggregated proteins in the affinity-purified GSS proteins by PROTEOSTAT staining. The experiment was performed with 3 technical replicates. **(C)** Venn diagram representing the number of unique semi-tryptic peptides ending with N identified in the insoluble fraction or whole cell lysate of HEK293T and MOLM-13 cells. **(D)** Quantification of percentage of unique semi-tryptic peptides ending with N out of all peptides identified in the insoluble fraction or whole cell lysate of HEK293T and MOLM-13 cells. **(E)** Volcano plots of semi-tryptic peptides ending with N in the insoluble fractions of cells after genetic knockout of CRBN compared to wild-type for HEK293T and MOLM-13 cells. The number of upregulated and downregulated peptides are noted. The experiment was performed with 4 technical replicates. P-values for the abundance ratios were calculated by one-way ANOVA with TukeyHSD post-hoc test.

To probe the connection between the cleavage events and protein aggregation in cells more broadly, we isolated the proteins that were insoluble in detergent (INSOL) from the whole cell lysate (WCL) using an adapted differential centrifugation method(62). The proteomic profiling of the WCL and INSOL fractions from HEK293T or MOLM-13 cells, an acute myeloid leukemia cell line used as a model to study the efficacy of CRBN-targeting agents, uncovered many cleavage sites unique to the isolated insoluble fractions (**Fig. 7C, Fig. S7C**). Notably, the insoluble fractions had a higher proportion of peptides cleaved at N or Q out of all the identified peptides in both cell lines, with a larger difference observed in HEK293T (**Fig. 7D, Fig. S7D**), suggesting that the cleavage products are prone to aggregation. Moreover, TMT-based quantitative proteomics of the insoluble fractions of WT and CRBN-KO cells revealed that the cleavage peptides were enriched upon loss of CRBN for HEK293T relative to the total protein level, although this effect is less pronounced for MOLM-13 cells (**Fig. 7E, Fig. S7E–F, Dataset S9**). These data indicate that the ability of CRBN to intercept cleaved fragments may be important to prevent their formation of otherwise insoluble aggregates and sheds light on the role of CRBN in quality control and maintenance of protein homeostasis.

## Discussion

We utilized proteomic, computational, and chemical biology approaches to characterize the spontaneous genesis of C-terminal cyclic imide modifications from aging or stressors. These data show that the intramolecular cleavage process is intrinsic to primary sequence, influenced by secondary structure, and accelerated by external stressors, like oxidation, heat, or elevated pH. The consensus sequence and structural features of C-terminal cyclic imide sites inform the proteins and residues susceptible to truncation, as well as hotspots of aggregation and other unwanted biological consequences. The trends observed from the meta-analysis are broadly reflected by experiments, confirming the susceptibility of sequences with small, polar residues at the +1 position and stability of those with hydrophobic, bulky residues at the +1 position. These experimental observations are expected for nucleophilic addition chemistry, which requires facile proton transfer and is hindered by sterics, and also align with the necessity of side chain H-bond capability for the side reaction of chemical protein synthesis that leads to cN formation via intramolecular cleavage(63). H or S at the +1 position also promotes asparagine deamidation(37), whereas asparagine with P at the +1 position cannot go through the deamidation pathway, which may rationalize the enrichment of NPK in the meta-analysis consensus sequence for cN events despite the faster cN formation rate of NHK or NSK sequence measured by experiments.

The intramolecular cleavage events were also accelerated by buffer components, heat and high pH, which indicates the possible upregulation of the modifications in specific developmental or disease states where the cellular environment undergoes a shift in these external factors(64–66). These factors are common themes that promote other protein damage modifications, implying that quality control mechanisms via CRBN participate in a broader protein damage recognition and removal process that is responsive to the elevation of protein damage events upon stress. The cleavage half-lives for the asparagine-bearing sequences, ranging from 100 to 2000 days for the species and conditions evaluated in this study, demonstrate that this form of protein damage can elicit biological responses within the time scale of many life forms. However, the requirement of detecting these species by mass spectrometry remains a challenge to readily measuring the extent of these modifications across sample types. Future development of selective enrichment methods for C-terminal cyclic imides will enable more rapid and sensitive identification of these modifications across different biological samples to probe the response of these modifications to different perturbations and complement the insights from proteomic profiling.

The observation that asparagine residues form C-terminal cyclic imides much faster than glutamine, together with the higher affinity of C-terminal aspartimides towards CRBN measured by TR-FRET(50), supports the hypothesis that CRBN preferentially recognizes cN as the more biologically prevalent modification. A systematic study on the substrate scope and preference of sequence recognition will further unveil the native substrates of CRBN and implications of the C-terminal cyclic imide modification beyond GSS and its aggregation. The dysregulation of the cleavage products may contribute to the association of the mutants of CRBN with neurological disorders, as seen in the link between PCMT1 loss and significant accumulation of isoaspartate protein damage in the brain. While we have surveyed the occurrence and spontaneous genesis of C-terminal cyclic imides, identification of alternate mechanisms of C-terminal cyclic imide formation through other non-enzymatic or enzymatic mechanisms may open new avenues for identification of additional CRBN substrates, synthetic generation of diverse CRBN ligands, and interrogation of biological pathways involving C-terminal cyclic imides in a CRBN-dependent or -independent manner. Exploration of E3 ligase adapters or repair proteins that regulate other non-enzymatic post-translational modifications is another area of future investigation that will extend our understanding of the biology of protein damage. In sum, the elucidation of the features that promote the C-terminal cyclic imide form of protein damage and their regulation in the presence or absence of CRBN provides a comprehensive model of how cells combat the inevitable damage to the proteome over the course of aging.

**Supplementary Information** and **Datasets** are available for this paper.

## Data Availability

Proteomics data have been deposited to the PRIDE repository with the dataset identifier PXD054016. All codes used in this study are available in GitHub: https://github.com/christinawoo/protein_damage_code. All study data are included in the main manuscript, SI Appendix, and Datasets S1–S9.

## Author Contributions

C.M.W. and W.X. conceptualized the study. W.X., N.C., and A.J. designed and performed peptide mass spectrometry experiments. W.X. performed in vitro, cellular, and proteomic studies and the analysis of data. M.S. and W.X. performed the computational analysis. Z.Z. and A.J. designed and synthesized compounds. W.X., H.C.L., and Z.Z. generated the proteins used in this study. E.Y.F. performed electroporation experiments. W.X. drafted the manuscript, and all authors reviewed and edited the manuscript.

## Competing Interest Statement

The authors declare the following competing interests: Harvard University has filed a PCT patent application on April 13, 2022 covering the chemical structures and their use. C.M.W. and W.X. are inventors of this patent. The Woo lab receives or has received support from Merck and Ono Pharmaceuticals. All other authors declare no competing interests.

## Acknowledgments

We thank S. Ichikawa, M. Anthony Leon-Duque, L. Yang, A. Scott, and Y. Li for helpful discussions and M. Chen and S. Trager from the Harvard University Mass Spectrometry and Proteomics Resource Laboratory for technical support. His_6_-CRBN/DDB1 was a generous gift from Boehringer Ingelheim. Some data used in this publication were generated by the Clinical Proteomic Tumor Analysis Consortium (NCI/NIH). Support from the Blavatnik Biomedical Accelerator at Harvard University (C.M.W.) and the National Institutes of Health (R01GM141406, C.M.W.) is gratefully acknowledged.

