## Supplementary Information for "Genesis and regulation of C-terminal cyclic imides from protein damage"

**Affiliations:**
† Department of Chemistry and Chemical Biology, Harvard University, Cambridge, MA 02138
### Novo Nordisk Research Center Seattle, Inc., Seattle, WA 90089

**Corresponding author:** Christina M. Woo

**This PDF file includes:**

Figures S1 to S11
Methods
Materials and Instrumentation
Synthetic Procedures
LC-MS traces
Legends for Datasets S1 to S9
SI References

**Other supplementary materials for this manuscript include the following:**

Datasets S1 to S9

**Table of Contents**


| | | |
|---|---|---|
| I. | Figures S1–S11 | S1 |
| II. | Methods | S13 |
| II. | Materials and Instrumentation | S25 |
| IV. | Synthetic Procedures | S31 |
| V. | Catalog of LC-MS traces | S32 |
| VI. | Legends for Datasets S1–S9 | S35 |
| VII. | SI References | S36 |

#### I. Supplementary Figures

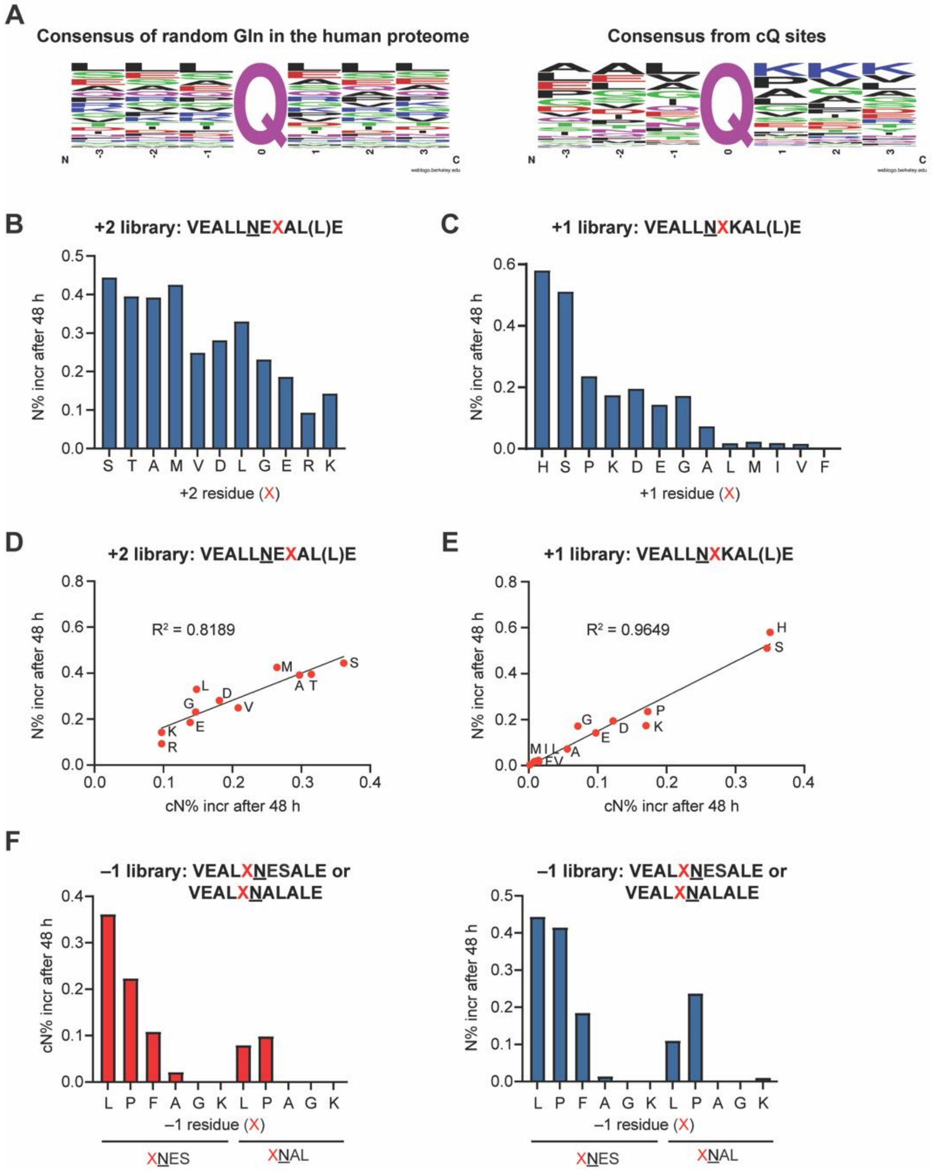


**Figure. S1. Investigation of the sequence dependence of C-terminal cyclic imide formation.** **(A)** Frequency charts of the sequence alignment spanning the –3 to +3 residues flanking random glutamine sites extracted from the human proteome and C-terminal glutarimide sites observed in CPTAC datasets. Alignments were generated with weblogo.berkeley.edu. **(B–C)** Quantification of N formation from +2 **(B)** or +1 **(C)** peptide library after 48 h incubation in 100 mM $Na_2HPO_4$ (pH 7.4) at 37 °C. **(D–E)** Correlation between cN and N formation for +2 **(D)** or +1 **(E)** peptide library after 48 h incubation in 100 mM $Na_2HPO_4$ (pH 7.4) at 37 °C. The coefficients of determination $R^2$ for the simple linear regression are noted. **(F)** Quantification of cN and N formation from –1 peptide library after 48 h incubation in 100 mM $Na_2HPO_4$ (pH 7.4) at 37 °C.

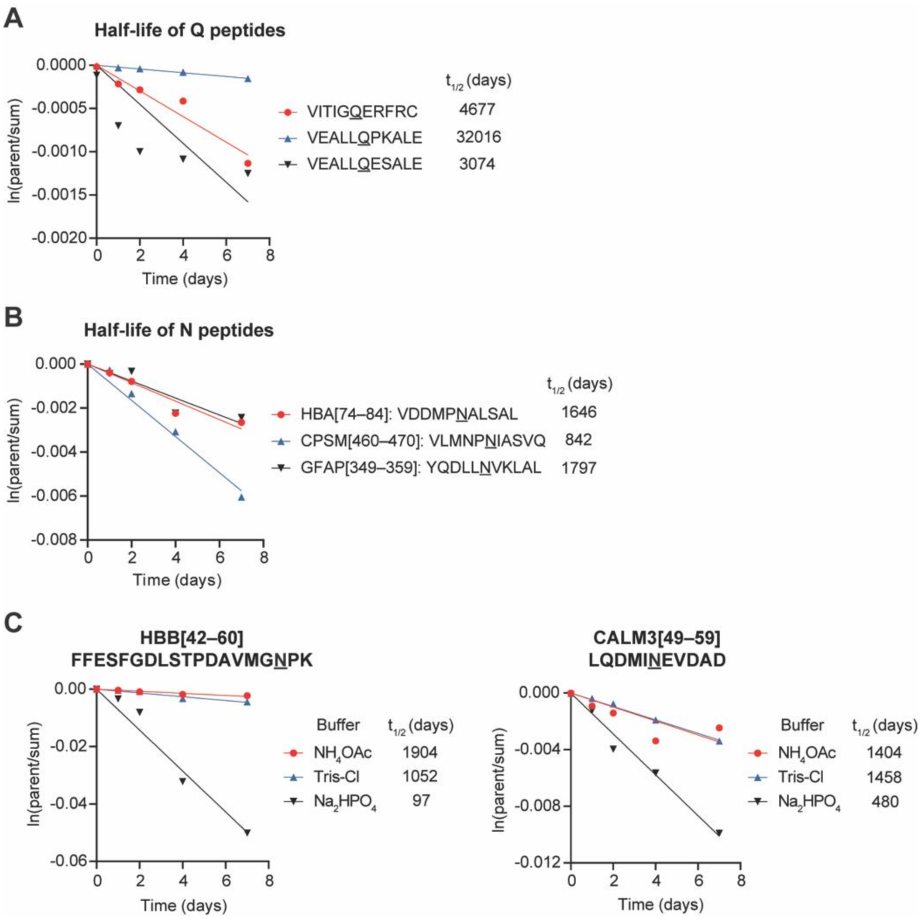


**Figure. S2. Characterization of intramolecular cleavage on representative peptide sequences.** **(A)** Time-course cleavage of three Q peptides in 100 mM $Na_2HPO_4$ (pH 7.4) at 37 °C. **(B)** Time-course cleavage of three stable peptides selected from frequent cN sites in the proteome in 100 mM $Na_2HPO_4$ (pH 7.4) at 37 °C. **(C)** Time-course cleavage of the indicated susceptible peptides in 100 mM $NH_4OAc$, Tris-Cl or $Na_2HPO_4$ (all pH 7.4) at 37 °C.

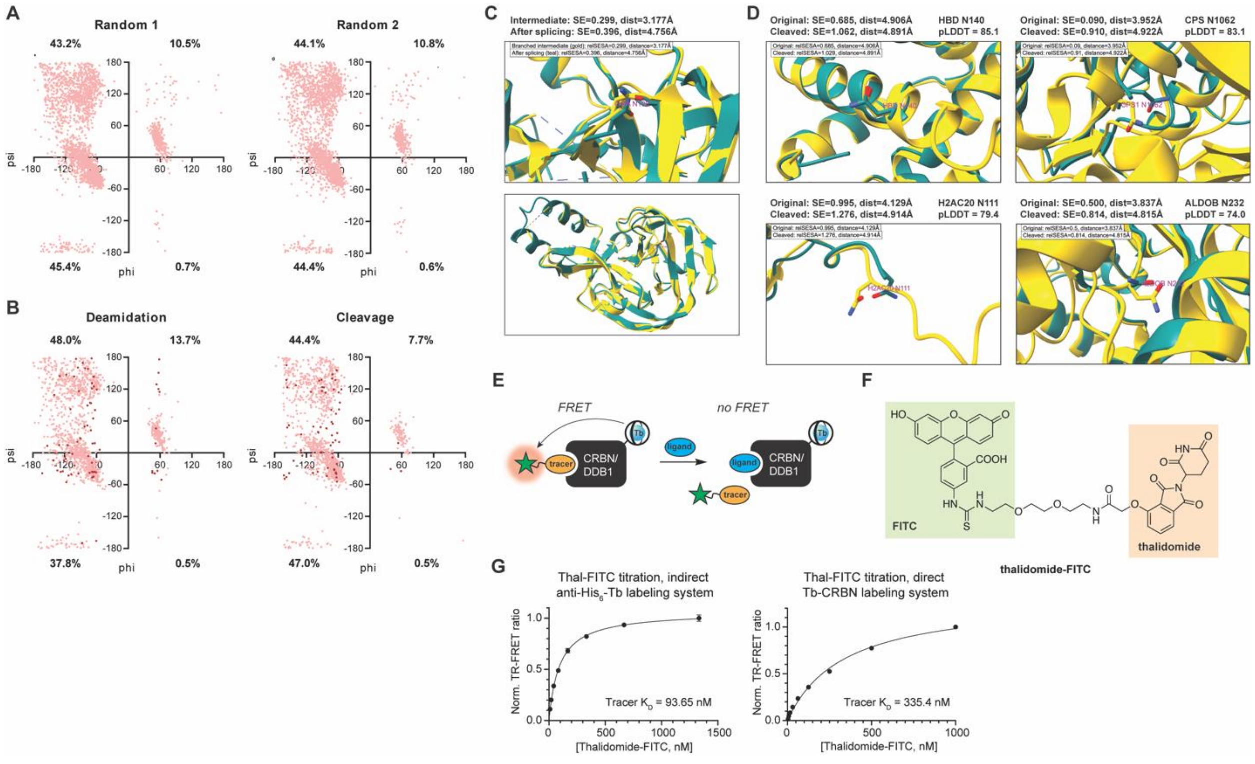


**Figure. S3. Structural features for the frequent cleavage sites. (A–B)** Ramachandran plots of two background datasets comprised of randomly extracted Asn sites **(A)** and the top frequent deamidation and intramolecular cleavage sites **(B)**. **(C)** Overlay of experimental structures of the branched intermediate of the Mxe GyrA intein in gold (PDB: 4OZ6) and the intein after splicing in teal (PDB: 1AM2). The calculated values of surface exposure and distance are noted. **(D)** Overlay of the original experimental structure in yellow and the predicted structure after cleavage at the indicated asparagine residue in teal for HBD N140, CPS N1062, H2AC20 N111, and ALDOB N252. The calculated values of surface exposure and distance and the pLDDT scores for the prediction of asparagine cleavage sites are noted. **(E)** Schematic of the TR-FRET assay used to measure the binding of ligands based on their ability to displace the tracer molecule. **(F)** Structure of thalidomide-FITC, the tracer used to assess the engagement with $His_6$-CRBN/DDB1 in this study. **(G)** Saturation binding of thal-FITC to $His_6$-CRBN/DDB1 labeled with CoraFluor-1-anti-$His_6$ conjugates or directly labeled with CoraFluor-1 with the determined $K_D$ values noted. All TR-FRET experiments were performed with 3 technical replicates.

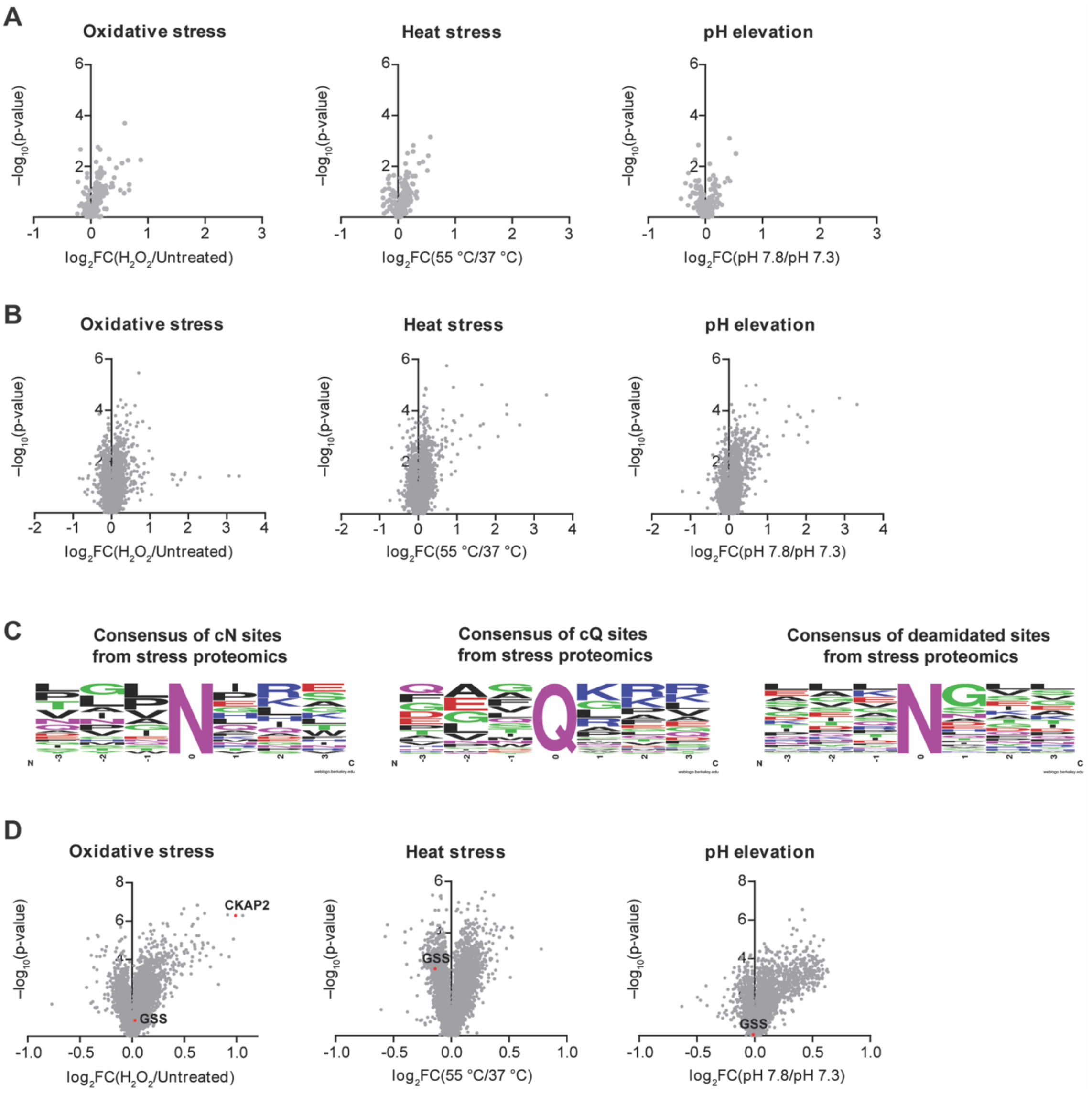


**Figure. S4. Proteomic profiling of stress-induced cleavage events in cells. (A–B)** Volcano plots of semi-tryptic peptides ending with Q **(A)** or fully tryptic peptides bearing asparagine deamidation modifications **(B)** in CRBN-KO HEK293T treated with 100 µM $H_2O_2$ for 40 min, incubated at 55 °C for 40 min, or incubated at pH 7.8 for 4 h compared to untreated cells incubated at pH 7.3, 37 °C. P-values for the abundance ratios were calculated by one-way ANOVA with TukeyHSD post-hoc test. **(C)** Frequency charts of the sequence alignment spanning the –3 to +3 residues flanking cN, cQ or asparagine deamidation sites identified from this proteomics experiment. Alignments were generated with weblogo.berkeley.edu. **(D)** Volcano plots of high-confidence proteins in CRBN-KO HEK293T treated with 100 µM $H_2O_2$ for 40 min, incubated at 55 °C for 40 min, or incubated at pH 7.8 for 4 h compared to untreated cells incubated at pH 7.3, 37 °C. The experiment was performed with 4 biological replicates. P-values for all the abundance ratios were calculated by one-way ANOVA with TukeyHSD post-hoc test.

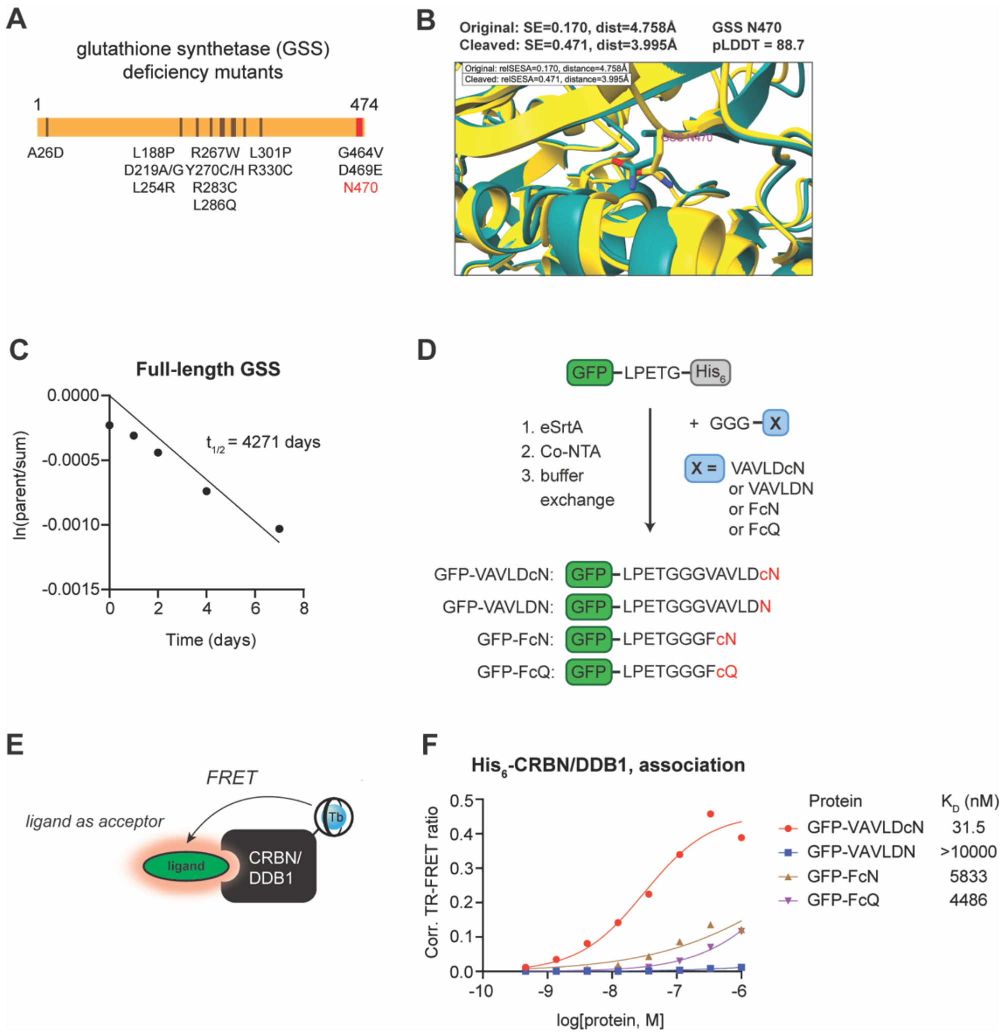


**Figure. S5. CRBN recognizes the GSS cN470 sequence. (A)** Schematic of mutants associated with GSS deficiency on the native sequence of human GSS (hGSS). **(B)** Overlay of the original experimental structure in yellow and the predicted structure after cleavage at the indicated asparagine residue in teal for GSS N470. The calculated values of surface exposure and distance and the pLDDT score for the prediction of asparagine cleavage site are noted. **(C)** Derivation of intramolecular cleavage half-life for full-length GSS protein in 100 mM $Na_2HPO_4$ (pH 7.4) at 37 °C. **(D)** Schematic of sortase system used to generate the tagged GFP used in this study. **(E)** Schematic of the TR-FRET assay used to measure the direct association of fluorescent ligands with $His_6$-CRBN/DDB1 complex. **(F)** Dose titration of the indicated tagged GFP in TR-FRET assay with the determined $K_D$ values noted. The experiment was performed with 3 technical replicates.

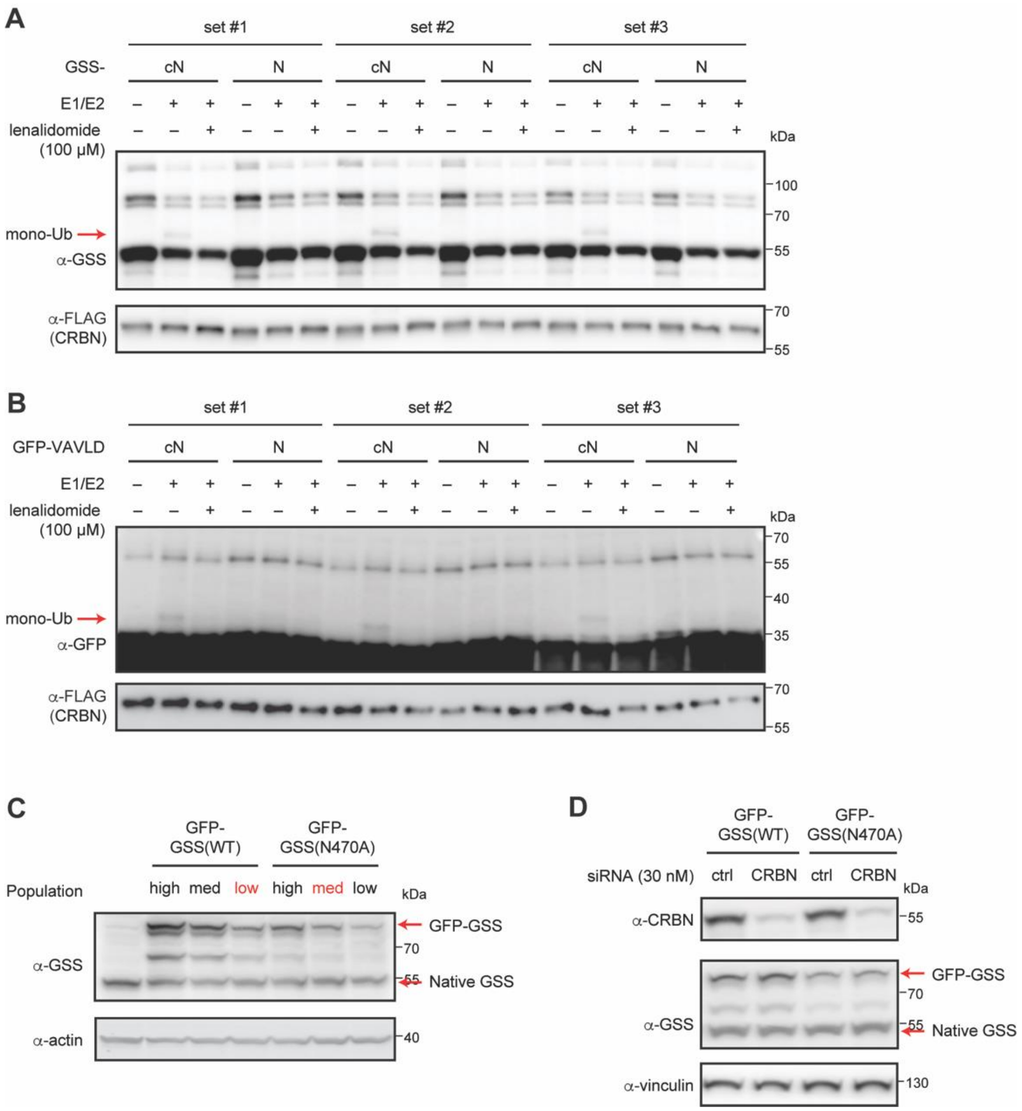


**Figure. S6. CRBN regulates GSS cN470 in vitro and in cells. (A)** In vitro ubiquitination of GSS-cN or GSS-N with K0 ubiquitin across three technical replicates. **(B)** In vitro ubiquitination of GFP-VAVLDcN or GFP-VAVLDN with K0 ubiquitin across three technical replicates. **(C)** Western blot analysis of GSS reporter cell lines sorted based on different GFP expression level. "Low" population from WT and "med" population from N470A were selected for further experiments due to their similar expression of GFP-GSS fusion protein. **(D)** Western blot analysis of GSS reporter cell lines transfected with 30 nM non-targeting siRNA (sictrl) or siRNA targeting CRBN (siCRBN) and incubated for 5 days post-transfection.

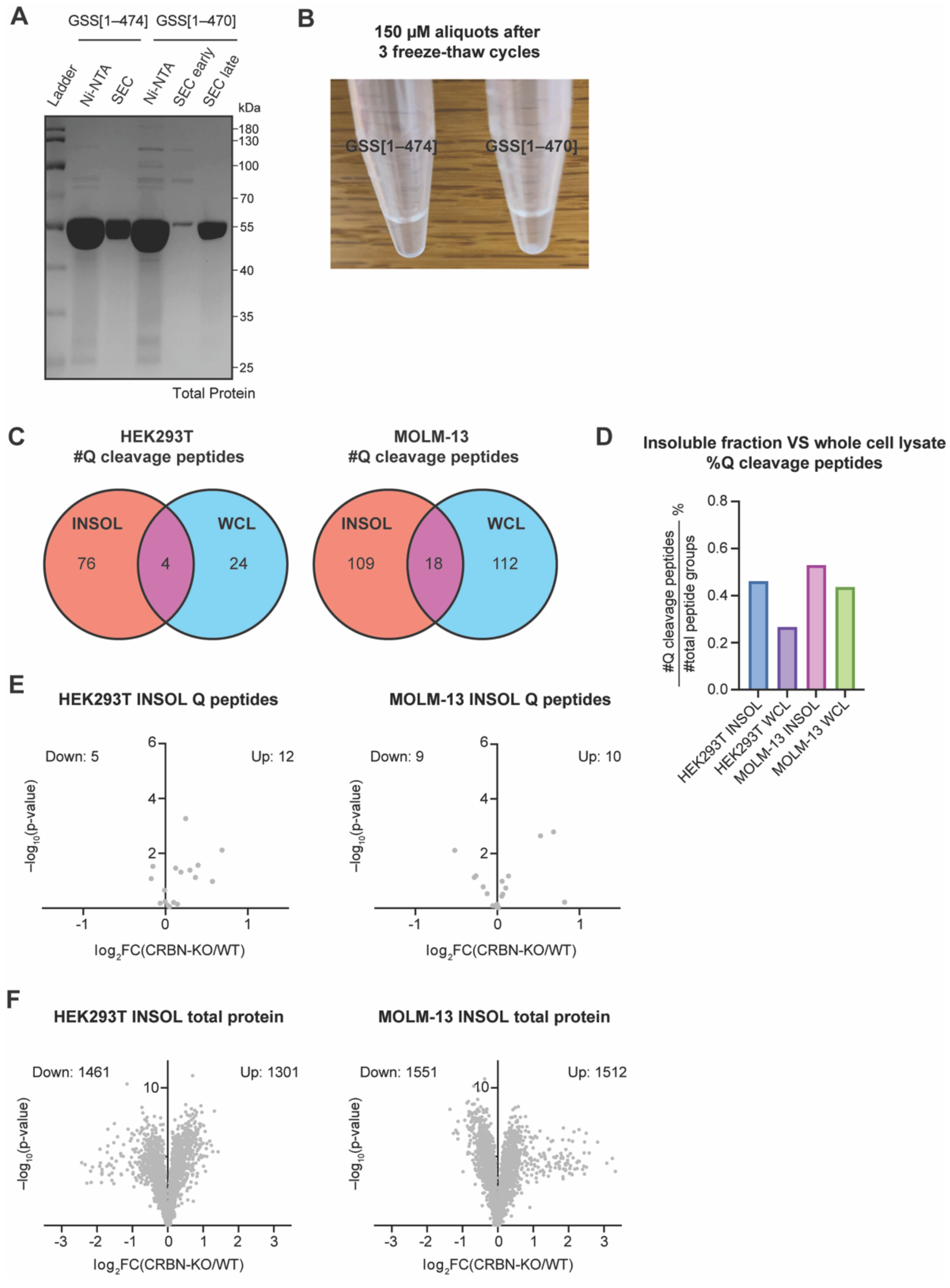


**Figure. S7. Cleavage of GSS and cellular proteins correlates with aggregation. (A)** SDS-PAGE analysis of the indicated GSS constructs. **(B)** Comparison of 150 µM aliquots of full-length and cleaved GSS after three freeze-thaw cycles. Apparent precipitation was observed for cleaved GSS. **(C)** Venn diagram representing the number of unique semi-tryptic peptides ending with Q identified in the insoluble fraction or whole cell lysate of HEK293T and MOLM-13 cells. **(D)** Quantification of percentage of unique semi-tryptic peptides ending with Q out of all peptides identified in the insoluble fraction or whole cell lysate of HEK293T and MOLM-13 cells. **(E)**

Volcano plots of semi-tryptic peptides ending with Q in the insoluble fractions of cells after genetic knockout of CRBN compared to wild-type for HEK293T and MOLM-13. The number of upregulated and downregulated peptides are noted. **(F)** Volcano plots of high-confidence proteins in the insoluble fractions of cells after genetic knockout of CRBN compared to wild-type for HEK293T and MOLM-13. The number of upregulated and downregulated proteins are noted. The experiment was performed with 4 technical replicates. P-values for the abundance ratios were calculated by one-way ANOVA with TukeyHSD post-hoc test.

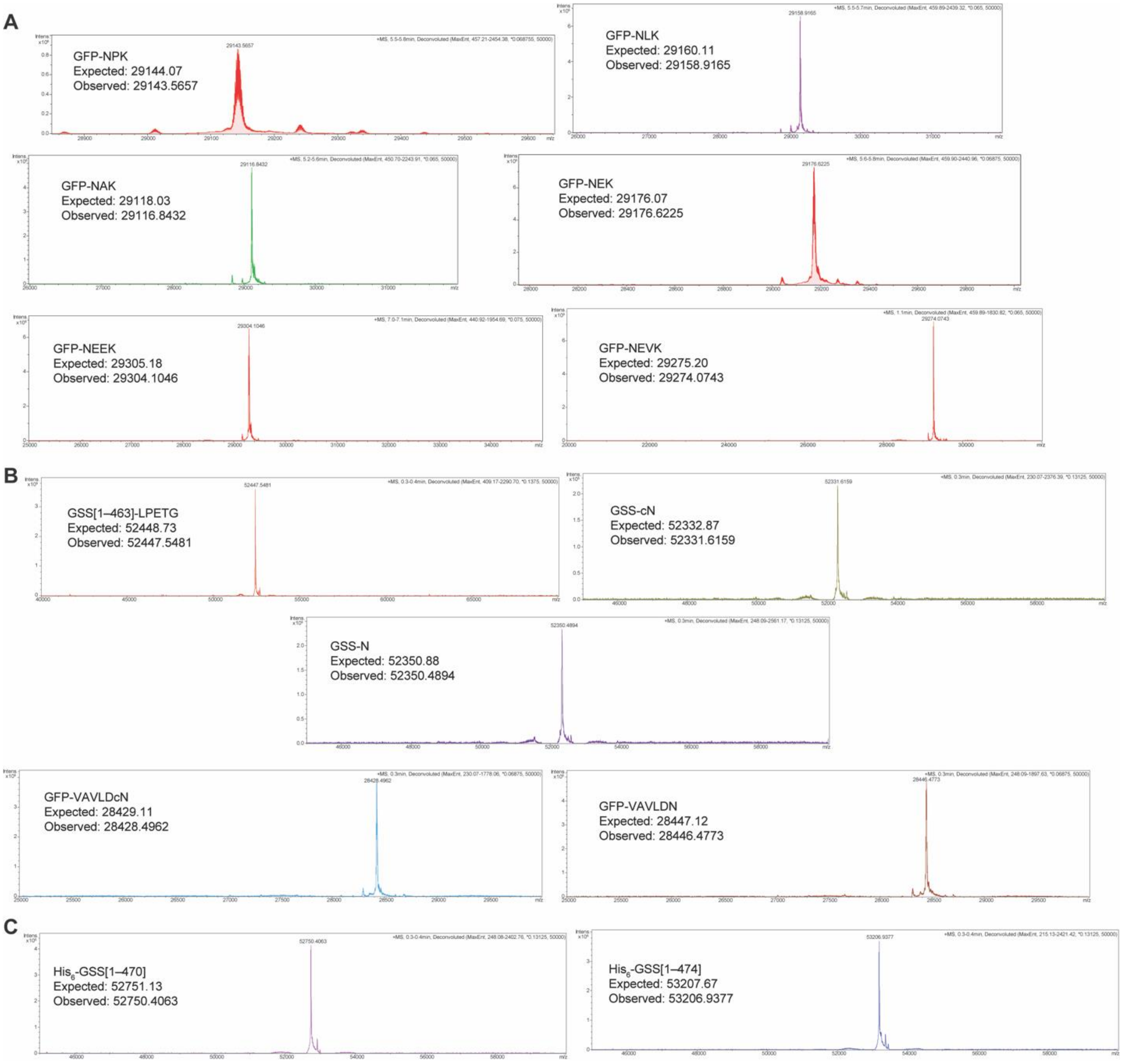


**Figure. S8. Characterization of recombinant proteins used in this study. (A)** Intact MS spectra of GFP model proteins. **(B)** Intact MS spectra of sortase reacted proteins. **(C)** Intact MS spectra of His$_6$-GSS[1–470] and His$_6$-GSS[1–474].

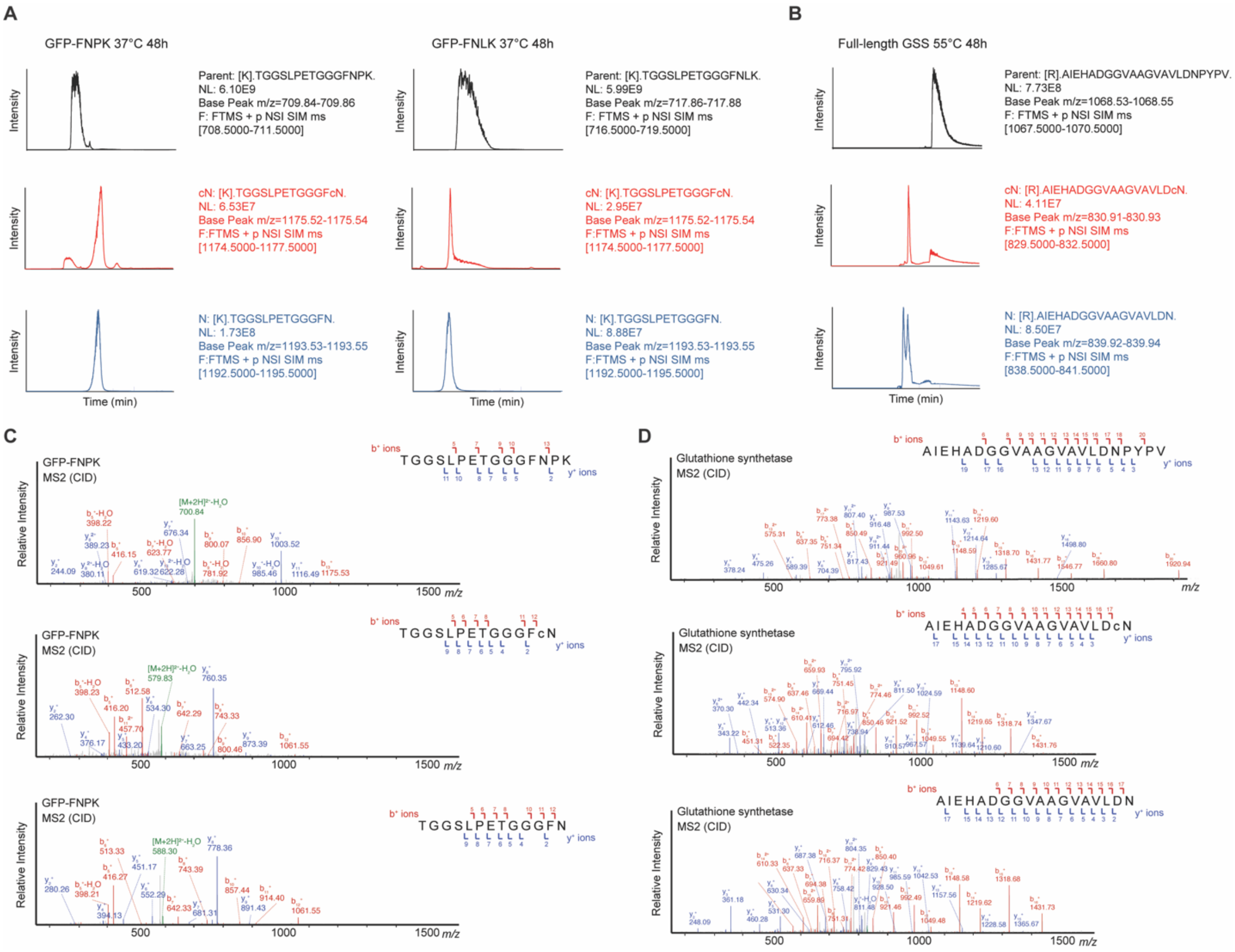


**Figure. S9. Representative chromatograms and MS2 spectra of the modified peptides identified from digested proteins. (A)** Example extracted ion chromatograms of parent, cN and N peptide ions from GFP model proteins. **(B)** Example extracted ion chromatograms of parent, cN and N peptide ions from full-length GSS protein. **(C)** Example MS2 spectra of parent, cN and N peptide fragments from GFP model protein. **(D)** Example MS2 spectra of parent, cN and N peptide fragments from full-length GSS protein.

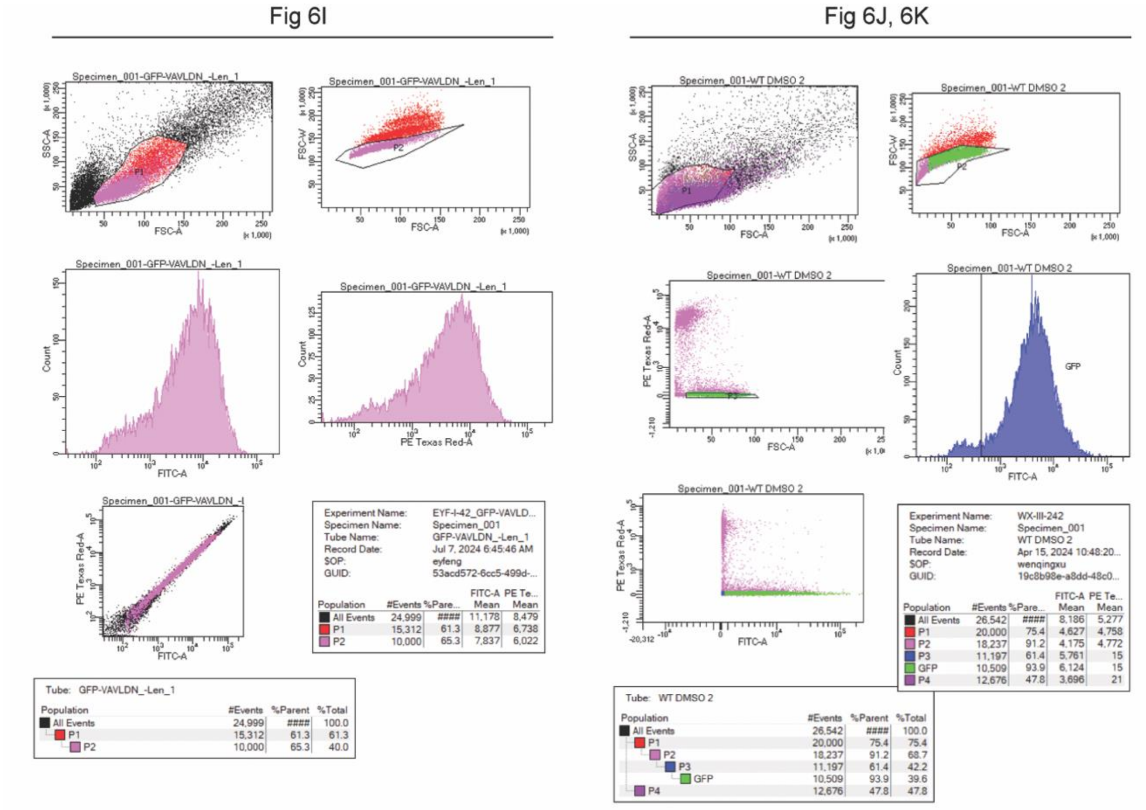


**Figure. S10. Representative flow cytometry gating strategy for Figure 6.**

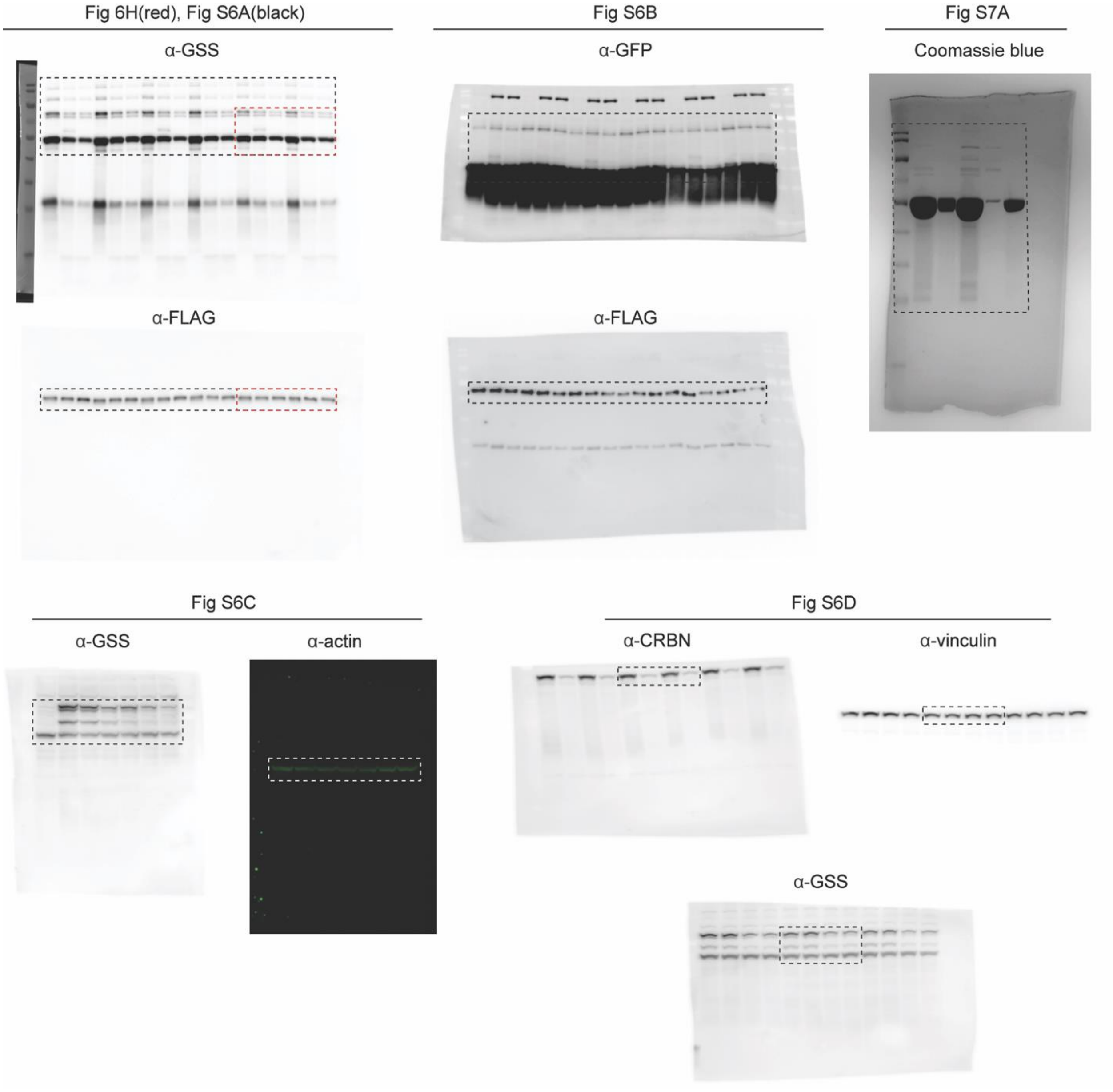


**Figure. S11. Uncropped western blot and gel images.**

#### II. Methods

**Formation study of C-terminal cyclic imide and its hydrolysis products in synthetic peptides.** The synthetic peptides were dissolved in $ddH_2O$ and incubated in 100 mM $Na_2HPO_4$, $NH_4OAc$, or Tris-HCl buffer at the indicated pH (final peptide concentration: 0.5 or 1 mM). Samples were incubated in a sand bath pre-heated to 37 °C, and an aliquot was taken at t = 0 h (collected immediately after resuspension in buffer) and the desired time points. The samples were immediately moved to –20 °C and stored until analysis. The MS samples were prepared by mixing 1 µL of the incubation mixture and 19 µL 0.1% formic acid. Samples (2 µL) were injected on a Thermo Orbitrap Fusion Lumos Tribrid and eluted using a multi-step gradient at a flow rate of 0.3 µL/min over 90 min (0−10 min, 5% acetonitrile in 0.1% formic acid/water; 10−72 min, 5−32% or 5−45%; 72−80 min, 32−98% or 45−98%; 80−90 min, 98%). The electrospray ionization voltage was set to 2 kV and the capillary temperature was set to 275 °C. MS1 scans were performed over 300–2000 or 350–2000 m/z at resolution of 120,000. The raw chromatograms were extracted with m/z of label-free parent peptide, cyclic imide, and hydrolysis products. The list of EIC mass ranges for the evaluated species can be found in Dataset S3, S4 and S7. For each species, only the peak area of the lightest isotope at the expected retention time was integrated and quantified using Xcalibur Qual Browser version 3.0.63. The two constitutional isomers of hydrolysis products were not distinguished in this study. Assuming the internal cleavage follows first-order kinetics for the analyzed time points, the simple linear regression of ln[(parent)/(parent+cN+N)] values as a function of time (days) was graphed by Prism 10 with constraint of the fitted line going through point (0, 0). The rate constant k was defined as the absolute value of the slope. The half-life of internal cleavage $t_{1/2}$ for each parent peptide was calculated using the last equation below.

$$\ln[parent]_t = \ln\left[\frac{parent}{parent + cN + N}\right]$$

$$\ln\frac{[parent]_t}{[parent]_0} = -\mathrm{kt}$$

$$t_{1/2} = \frac{ln(1/2)}{-\mathrm{k}} \approx \frac{0.693}{k}$$

**Generation of GFP model proteins.** pET28a-GFP-LPETGGGFNPK (plasmid 2) was first constructed from pET28a-GFP-LPETG-$His_6$ by adding the sequence encoding GGFNPK before the $His_6$ tag with primers 1–2 and Q5 site-directed mutagenesis kit (NEB E0554S). The desired +1 position mutants (plasmids 3–5) were generated from plasmid 2 with the corresponding overlapping primers (primers 3–8) and QuikChange Lightning kit (Agilent 210518). For the +2 position mutants, plasmid 6 was constructed from plasmid 2 using primers 9–10 with Q5 site-directed mutagenesis kit and then plasmid 7 was generated from plasmid 6 using primers 11–12 with QuikChange Lightning kit. All plasmids were sequence verified by Sanger sequencing before use. The overexpression and purification of GFP proteins were performed as previously reported(1). In brief, BL21(DE3) cells were transformed with the plasmid and used to inoculate overnight cultures of LB + 50 µg/mL kanamycin. Large-scale overexpression cultures (750 mL×2 for each mutant) were inoculated with overnight culture diluted 1:100. The overexpression cultures were incubated at 37 °C with shaking at 200 rpm until the OD600 was approximately 0.6, at which point the temperature was reduced to 20 °C and IPTG was added to a final concentration of 0.45

mM. The cultures were incubated for approximately 16 h prior to collecting the cells by centrifugation. To purify protein, cell pellets were resuspended in 8.3 mL of 20 mM imidazole, 1× protease inhibitor, 1% Triton-X 100/PBS per pellet. Lysates were sonicated (30 sec on, 10 sec off, 5 min total, 25% amplitude) on ice and clarified by centrifugation (20,000 × g, 4 °C, 10 min) and syringe filtration (0.45 μm). The His-tagged protein was first purified on a Ni-bound 1 mL HiTrap Chelating HP column, equilibrating and washing with 20 mM imidazole/PBS and eluting with a gradient to 500 mM imidazole/PBS. Protein-containing fractions were concentrated to < 1 mL and further purified on a S75 10/300 GL column, pre-equilibrated and run with 100 mM ammonium bicarbonate (pH 7.4). Protein-containing fractions were collected and concentrated with a 10 kDa MWCO spin filter, and the concentration determined by A488 (ε = 55,000 $M^{-1}$ $cm^{-1}$). The protein was aliquoted, lyophilized, and stored at –80 °C.

**Intact protein mass spectrometry.** An Agilent PLRP-S (50 mm length, 5 μm particle size, 4.6 mm ID, 1000 Å pore size) was used. The column was maintained at 70 °C during the run. For each sample, 10 μL protein solution was either injected directly through a union and eluted with 0.1% formic acid/60% acetonitrile/water or injected onto the column and eluted using the following method with mobile phases A (0.1% formic acid/water) and B (0.1% formic acid/acetonitrile). Prior to the gradient, the column was maintained at 0% B for 2 min to wash salts. Then, a linear gradient was applied over 10 min to a final concentration of 100% B. The column was maintained at 100% B for 1 min to wash before changing to 0% B over 0.1 min. Then, the column was re-equilibrated at 0% B for 5.9 min prior to the next run. The data was internally calibrated using sodium formate clusters injected at the end of each run. The data was analyzed using Bruker Compass DataAnalysis (v.4.3), and deconvoluted using the maximum entropy algorithm between selected mass ranges. Modification masses used included: GFP fluorophore formation = –20.77 Da; N-terminal methionine loss = –131.20 Da.

**Formation study of C-terminal cyclic imide and its hydrolysis products in recombinant proteins.** GFP model proteins (75 μg) were aliquoted and lyophilized per condition resuspended in 100 mM $Na_2HPO_4$ at the indicated pH. GSS protein (100 μg) was aliquoted from the commercial tube and buffer exchanged into 100 mM $Na_2HPO_4$ at the indicated pH with Zeba columns. Proteins were incubated in a sand bath pre-heated to varying temperatures. An aliquot from each sample was collected at t = 0 h (collected immediately after resuspension) and the desired time points. Immediately after collection, the samples were stored at –80 °C until analysis.

All the samples were dried on a vacufuge at once and resuspended in 23 μL of 5% SDS, 50 mM TEAB, pH 7.55. The samples were reduced by addition of dithiothreitol (1 μL of 240 mM stock) at 24 °C for 30 min then alkylated by addition of iodoacetamide (1 μL of 500 mM stock) and incubation in the dark at 24 °C for 30 min. The samples were acidified by the addition of phosphoric acid (2.5 μL of 27.5% stock) to a final concentration of 2.5%. S-Trap buffer (90% methanol, 0.1 M TEAB, pH 7.1, 165 μL) was then added. Each sample was transferred to a S-Trap micro column. Using a vacuum manifold, the columns were washed with S-Trap buffer (3 × 180 μL). To digest the S-trap-bound proteins, 2 μg of trypsin resuspended in 20 μL 50 mM TEAB pH 7.4 was added to each column and incubated at 47 °C for 1 h without agitation. The digested peptides were eluted by sequential addition of 50 mM TEAB pH 7.4 (40 μL), dd$H_2O$ (40 μL) and 0.2% formic acid, 50% acetonitrile/water (40 μL), with each elution collected by centrifugation (4,000 × g, 24 °C, 1 min) in a clean Eppendorf tube. The dried peptide digests were fractionated into 5 fractions using the Pierce high pH reversed-phase peptide fractionation kit. The peptides were eluted sequentially by 5% (excluded from analysis), 10%, 15%, 20%, 35% and 50%

acetonitrile/0.1% TEA. Immediately after the elution, each fraction was acidified by the addition of 5 µL of 10% formic acid. The fractions were concentrated to dryness and each sample was resuspended in 20 µL of 0.1% formic acid. The samples were injected on a Thermo Orbitrap Fusion Lumos Tribrid and eluted using a multi-step gradient at a flow rate of 0.3 µL/min over 90 min (0−10 min, 5% acetonitrile in 0.1% formic acid/water; 10−72 min, 5−32%; 72−80 min, 32−98%; 80−90 min, 98%). The electrospray ionization voltage was set to 2 kV and the capillary temperature was set to 275 °C. MS1 scans were performed on the selected ion monitoring (SIM) mode to only look for narrow windows of m/z for the parent peptide, cyclic imide and hydrolyzed products. Orbitrap resolution was set at 120,000, RF lens at 60%, maximum injection time at 300 ms and normalized AGC target at 2000%. The raw chromatograms were extracted with m/z of label-free parent peptide, cyclic imide, and hydrolysis products. For each protein, a test run with full-scan mode was initially conducted for all the fractions to search for the presence of the target species, and the fractions containing at least one of the species were subsequently analyzed by SIM, which were found to be fractions 1 and 2 (10% and 15% acetonitrile in 0.1% TEA) for all GFP mutants and GSS. The list of SIM m/z window and EIC mass range for all the evaluated peptides can be found in Dataset S3 and S7. For each species, only the peak area at the expected retention time was integrated and quantified using Xcalibur Qual Browser version 3.0.63. Half-life calculations were performed as described in "Formation study of C-terminal cyclic imide and its hydrolysis products in synthetic peptides".

**Identification of C-terminal cyclic imide and deamidation modification sites from public datasets.** The raw data for the global proteomics datasets were obtained from CPTAC and processed according to the methods in the respective publication for fully tryptic peptides and spectra that did not receive a confident peptide spectral match were searched for N-terminal semi-tryptic sequences and dynamic modification of dehydration on asparagine or glutamine residues (–18.015 Da) at the peptide C-terminus for internal cleavage and searched for fully tryptic sequences and dynamic modification of deamidation on asparagine and glutamine residues (+0.984 Da). The data were filtered with a 1% FDR using Percolator. The background datasets were generated by random extraction of 50,000 asparagine or glutamine sites from the Swiss-Prot human proteome. 7-mer sequence alignments for C-terminal cyclic imides, deamidation and random N/Q were generated from non-contaminant proteins with at least 3 amino acid residues following the modification sites in their intact sequences.

**Clustering of C-terminal cyclic asparagine modification sites.** The set of heptamer sequences representing –3 to +3 residues surrounding cN occurrences was analyzed using agglomerative hierarchical clustering. The distance between two individual samples was defined as $1 - f$, where $f$ is the fraction of identical amino acids in the +1 and +2 positions. This distance metric was used to identify clusters with high similarity in the +1 and +2 regions which yields a low average distance between sequences, using an average linkage approach. The clustering method was applied to partition the set of sequences into three clusters.

**Computational analysis of protein 3D structures.** The most frequent sites for deamidation and internal cleavage were compiled and aggregated using the pivot table based on the counts of the entry combining protein accession and residue position. The background datasets were generated by two rounds of random extraction of 5000 asparagine sites from the Swiss-Prot human proteome. For each site (deamidation, internal cleavage, and random), PDB structures containing the corresponding site and with resolution 3.5 Angstroms or finer were programmatically fetched from the Protein Data Bank in Europe (PDBe). If the structure containing the site and meeting the standard is unavailable, the structures predicted by AlphaFold(2) were fetched and further filtered to only those with pLDDT ≥ 70 for the target site. For each residue, the relative solvent-exposed surface area (relSESA), distance from side-chain N to backbone carbonyl C, and backbone torsion angles (psi/phi) were computed in UCSF ChimeraX using the fetched structures. According to the ChimeraX standard, relSESA value was calculated by computing the area of the solvent-excluded surface (command: *surface*) for the residue, then dividing by the solvent-excluded surface area of the corresponding residue in a Gly-X-Gly tripeptide(3). The distance from the side-chain N to backbone carbonyl C was calculated using the *distance* command, and backbone torsion angles were extracted from the residue attributes.

For selected sites of internal cleavage, the structure of the truncated sequence resulting from internal cleavage was predicted using the Google Colab version of AlphaFold. The predicted cleavage structure was then superimposed with the original full-length PDB structure in ChimeraX (command: *matchmaker*). The codes used in this study are available on GitHub: https://github.com/christinawoo/protein_damage_code.

**General cell culture protocol.** Human-derived cell lines were cultured in DMEM supplemented with 10% heat-inactivated fetal bovine serum (FBS) and 1× penicillin-streptomycin (noted as DMEM+/+). Cells were grown at 37 °C in a humidified atmosphere with 5% $CO_2$ unless otherwise noted. Mycoplasma testing was performed regularly for all cell lines to check for contamination. For the collection of cell pellets, cells were dissociated and collected by two PBS washes and centrifugation at 500 × g, 24 °C, 3 min. The pellets were flash frozen with liquid nitrogen and stored at –80 °C until use.

**Global quantitative proteomics sample preparation.** The proteomics sample preparation protocol was adapted from Donovan and coworkers(4) and off-line fractionation protocol was adapted from Batth and coworkers(5). Global proteomics samples were prepared in biological quadruplicate for each of the 4 conditions (untreated, oxidative stress, heat stress, pH elevation). $2.0\times10^6$ HEK293T CRBN-KO cells were seeded in 6-well plates and treated with each of the stress conditions for the indicated time: untreated cells were seeded in unadjusted DMEM+/+ (pH 7.3) and incubated at 37 °C for 4 h; for oxidative stress, cells were seeded in DMEM+/+ with 100 μM $H_2O_2$ for 40 min; for heat stress, cells were collected in a Falcon tube with DMEM+/+ and placed in a heat block preheated to 55 °C for 40 min; for pH elevation, cells were seeded in DMEM+/+ at pH 7.8 for 4 h. After treatment, the media was replaced with fresh, unadjusted DMEM+/+ and all samples were further incubated at 37 °C for a total of 24 h. Cells were collected by PBS washes, lysed by probe sonication (5 sec on, 3 sec off, 15 sec in total, 11% amplitude) in lysis buffer (8 M urea, 50 mM NaCl, 50 mM HEPES, 1×protease/phosphate inhibitor cocktail, pH 7.4), and cleared by centrifugation (21,000 × g, 4 °C, 10 min). After protein quantification by BCA protein assay, the lysates were diluted to 2 mg/mL with the lysis buffer. The samples (200 μg protein per sample) were reduced by addition of dithiothreitol (5 mM) at 24 °C for 30 min then alkylated by addition of iodoacetamide (15 mM) and incubation in the dark at 24 °C for 30 min. Proteins were precipitated by methanol-chloroform precipitation. Four volumes of methanol were added to the cell lysate, followed by one volume of chloroform, and finally three volumes of water. The mixture was vortexed and centrifuged at 14,000 × g for 5 min at 4 °C and the supernatant was carefully aspirated. The precipitated protein was washed with three volumes of methanol, centrifuged at 14,000 × g for 5 min at 4 °C, and the resulting protein pellet was dried for 10 min in a vacufuge. Precipitated protein was resuspended in 4 M urea, 50 mM HEPES pH 7.4 (25 μL) and 200 mM HEPES pH 7.4 (75 μL) was subsequently added for 1 M final urea concentration. The samples were first digested with LysC (2 μg) for 4 h at 22 °C. The mixture was then diluted by the addition of 200 mM HEPES pH 7.4 (100 μL), and further digested with trypsin (4 μg) for 16 h at 37°C. For each sample, half of the digested peptides (ca. 100 μg) was taken to labeling by TMTpro 16-plex reagent (34 μL). The labeling reactions were performed for 1.5 h at 22 °C and quenched by the addition of 1 M Tris-Cl, pH 7.6 (20 μL) and incubation for 15 min at 22 °C. The channels were combined, dried in a vacufuge, and desalted using a Pierce peptide desalting spin column (Thermo) according to the manufacturer's protocol. The desalted sample was offline fractionated into 80 fractions by high pH reverse-phase HPLC (Agilent 1260 Infinity II) through an Aeris peptide XB-C18 column (Phenomenex 00G-4507-E0) with mobile phase A containing 5% acetonitrile and 10 mM ammonium bicarbonate, and mobile phase B containing 90% acetonitrile and 1 mM ammonium bicarbonate (both pH 7.4). Fractions (0.7 mL each) were collected using a fraction collector in a deep 96-well plate. Samples were initially loaded onto the column at 1 mL/min for 4 min, after which the fractionation gradient was set as follows: 1% B to 27% B in 50 min, 60% B in 4 min, and ramped to 70% B in 2 min. Fraction collection was stopped at this point, and the gradient was held at 70% B for 5 min before being ramped back to 1% B to wash and equilibrate the column. The 80 resulting fractions were dried and pooled in a non-continuous manner into 20 fractions. The 20 final fractions were resuspended in 40 μL 0.1% formic acid.

**Proteomics mass spectrometry acquisition procedures.** The resuspended samples (2 μL) were run on an Orbitrap Eclipse Tribrid Mass Spectrometer coupled with a Vanquish Neo HPLC system (Thermo) at the Harvard Center for Mass Spectrometry. The peptides were first trapped on a trapping cartridge (300 μm x 5 mm PepMap™ Neo C18 Trap Cartridge, Thermo) prior to separation on a silica-chip-based micropillar column (μPAC, C18 pillar surface, 50 cm bed, Thermo). The column oven temperature was maintained at 35 °C. Peptides were eluted using a multi-step gradient at a flow rate of 300 nL/min over 180 min. The mobile phase A and weak wash liquid was water with 0.1% formic acid, and mobile phase B was acetonitrile with 0.1% formic acid, and strong wash liquid was 80% acetonitrile with 0.1% formic acid. The mobile phase gradient consists of a linear 125 min gradient from 2% to 20% mobile phase B, followed by a 24 min increase to 35% B, a further 10 min increase to 95% B, a 20 min plateau phase at 95% B. The autosampler temperature was 7 ºC. The column "Fast equilibration" was enabled. The Orbitrap Eclipse MS was operated in DDA mode with 3 s cycle time. The electrospray ionization was in positive mode with voltage of 2.1 kV and the capillary temperature at 275 ºC. Dynamic exclusion was enabled with a mass tolerance of 10 ppm and exclusion duration of 60 s. Full scan was performed in the range of 400–1,600 m/z at a resolution of 120,000, RF lens 30%, normalized AGC target 200% and maximum injection time set to auto. Precursors were isolated in a window of 0.7 m/z and charge states of 2–6. MS2 fragmentation was performed by HCD using normalized collision energy of 38% at the resolution of 50,000. The normalized AGC target was set to 250%, and a maximum ion injection time set to 200 ms.

**Mass spectrometry data analysis for protein level quantitation.** Analysis was performed in Thermo Scientific Proteome Discoverer version 2.4.1.15. The raw data were searched against SwissProt human (Homo sapiens) protein database (21 February 2019; 20,355 total entries) and contaminant proteins using the Sequest HT algorithm. Searches were performed with the following guidelines: spectra with a signal-to-noise ratio greater than 1.5; mass tolerance of 20 ppm for the precursor ions and 0.02 Da for the fragment ions; semi-trypsin digestion at the peptide N-terminus only to search for cleavage peptides and full trypsin digestion to search for deamidated peptides and quantify total protein level; fewer than 2 missed cleavages; static carboxyamidomethylation of cysteine residues (+57.021 Da); static TMTpro 16-plex labeling (+304.207 Da) at lysine residues and N-termini; variable oxidation on methionine residues (+15.995 Da); variable dehydration on asparagine and glutamine residues at C-terminus only (–18.015 Da). To search for deamidation sites, the variable dehydration modification was replaced with variable deamidation on asparagine residues (+0.984 Da). The TMT reporter ions were quantified using the Reporter Ions Quantifier node and normalized to the total peptide amount. Peptide spectral matches (PSMs) were filtered using a 1% false discovery rate (FDR) using Target Decoy PSM Validator. For the quantification of total proteins, the data were further filtered to include only master proteins with high protein FDR confidence, at least 3 unique peptides, and exclude all contaminant proteins. To derive the list of peptides generated from internal cleavage at N or Q, the data were filtered to include only peptides without oxidation modifications, bearing N or Q residues at the peptide C-terminus with or without dehydration, and derived from non-contaminant proteins. The peptides that were mapped back to the protein C-terminus were excluded. To derive the list of peptides with deamidation modifications, the data were filtered to include only peptides without oxidation modifications, bearing at least one deamidated N residue, no missed cleavages, and derived from non-contaminant proteins.  The abundance ratios and their associated p-values were calculated by one-way ANOVA with TukeyHSD post-hoc test.

**Overexpression and purification of recombinant GSS-LPETG.** pET28a-GSS[1–463]-LPETG-$His_6$ (plasmid 9) was constructed from the plasmids pcDNA3.1-FLAG-GSS (Genscript) and pET28a-GFP-LPETG-$His_6$(1). The insert corresponding to GSS[1–463] and backbone excluding GFP were amplified using overhang PCR with primers 13–16 and assembled using HiFi DNA Assembly Kit (NEB). The ligated product was transformed into NEB 5 alpha competent cells and the sequences validated by Sanger sequencing.

BL21(DE3) cells were transformed with the validated plasmid and used to inoculate overnight cultures of LB + 50 μg/mL kanamycin. Large-scale overexpression cultures (750 mL×4) were inoculated with overnight culture diluted 1:100. The overexpression cultures were incubated at 37 °C with shaking at 220 rpm until the OD600 was approximately 0.6, at which point the temperature was reduced to 16 °C and IPTG was added to a final concentration of 0.5 mM. The cultures were incubated for approximately 20 h prior to collecting the cells by centrifugation. To purify protein, cell pellets were resuspended in 8.3 mL of 20 mM imidazole, 1× protease inhibitor, 1% Triton-X 100/PBS per pellet. Lysates were sonicated (20 sec on, 10 sec off, 5 min total, 27% amplitude) on ice and clarified by centrifugation (20,000 × g, 4 °C, 10 min) and syringe filtration (0.45 μm). The His-tagged protein was first purified on a Ni-bound 1 mL HiTrap Chelating HP column, equilibrating and washing with 20 mM imidazole/PBS and eluting with a gradient to 500 mM imidazole/PBS (both pH 8.0). Protein-containing fractions were concentrated to < 1 mL and further purified on a S75 10/300 GL column, pre-equilibrated and run with PBS (pH 7.4). Protein-containing fractions were collected and concentrated with a 10 kDa MWCO spin filter, and the concentration determined by A280 (ε = 33,015 $M^{-1}$ $cm^{-1}$). Two separate species were detected on SEC, where the earlier fractions showed significant aggregation in PBS and thus the later fractions were stored and taken to further experiments.

**Sortase reaction.** For each reaction, 150 μL of 50 μM LPETG-containing protein was combined with 30 μL of 10 mM peptide substrate in 555 μL reaction buffer (100 mM Tris pH 7.5, 150 mM NaCl, 5 mM $CaCl_2$). Then, 15 μL of 97 μM eSrtA was added and the reaction was mixed by flicking. The reaction was allowed to incubate at 24 °C for 1 h. Non-reacted LPETG-containing protein and eSrtA were removed by adding 115 μL of washed His-purification Dynabeads and incubating with inversion for 15 min. The supernatant was collected using a magnetic tube rack.

The excess of peptide substrate was removed by buffer exchange into PBS with Zeba columns. The final concentration was determined by measuring the A280 for GSS (ε = 33,015 $M^{-1}$ $cm^{-1}$) or A488 for GFP (ε = 55,000 $M^{-1}$ $cm^{-1}$) and identity confirmed by intact protein MS.

**TR-FRET displacement assay to measure peptide affinity.** Anti-$His_6$ antibody (Abcam ab18184) was labeled with CoraFluor-1-Pfp ester as previously described to generate the TR-FRET donor. TR-FRET experiments were performed in Proxiplate-384 Plus (VWR PERK6008280) in 15 µL assay volume. TR-FRET measurements were acquired on a SpectraMax iD5 plate reader with SoftMax Pro software version 7.1.2, with the following settings: 350 nm excitation, 490 nm (Tb), and 520 nm (FITC) emission, 110 flashes per read, 0.05 ms excitation time, 0.05 ms delay and 0.2 ms integration. TR-FRET ratio was taken as the 520/490 nm intensity ratio per well.

To determine the tracer $K_D$ in the indirect labeling system, 5 nM CoraFluor-1-labeled anti-$His_6$ antibody and 10 nM $His_6$-CRBN/DDB1 complex were mixed in the assay buffer (25 mM HEPES, 150 mM NaCl, 0.5 mg/mL BSA, 0.005% TWEEN 20, pH 7.5). Thal-FITC was first added in serial dilution (1:2 titration, 8-point, $c_{max}$ = 1.33 µM) using an HP D300 digital dispenser and allowed to equilibrate for 10 min at room temperature before TR-FRET measurements were taken. The $K_D$ determined from the one-site model graphed in Prism 10 was used in the equation below to adjust for a two-site model due to the presence of bivalent anti-$His_6$ antibody:

$$K_{D,adj} = K_D \times (1 + \sqrt{2})$$

For ligand displacement assays, 5 nM CoraFluor-1-labeled anti-$His_6$ antibody,10 nM $His_6$-CRBN/DDB1 complex, and 50 nM Thal-FITC were mixed in the assay buffer. In all cases, small molecules or peptides were added in serial dilution (1:3 titration, 8-point, $c_{max}$ = 3 µM) using an HP D300 digital dispenser and allowed to equilibrate for 10 min at room temperature before TR-FRET measurements were taken. All experiments were performed in technical triplicates. Background signal (bottom) was determined from wells containing 100 µM 5-$NH_2$-lenalidomide to represent complete displacement. The assay ceiling (top) was defined via a no-ligand control. Data were background-subtracted, normalized and fitted to a four-parameter dose-response model [log(inhibitor) vs. response – Variable slope (four parameters)] in Prism 10, with constraints of Top = 1 and Bottom = 0 to derive the ligand $IC_{50}$ values. Ligand $K_D$ values have been calculated using Cheng-Prusoff principles, outlined in the equation below:

$$K_D = \frac{IC_{50}}{1 + \frac{[S]}{K_x}}$$

Where $IC_{50}$ is the measured $IC_{50}$ value, [S] is the concentration of fluorescent tracer, and $K_X$ is the adjusted $K_D$ of the fluorescent tracer.

**TR-FRET displacement assay to measure protein affinity.** The $His_6$-CRBN/DDB1 complex was labeled with CoraFluor-1-Pfp ester as previously described(6) to generate the TR-FRET donor. TR-FRET measurements were taken as described in the section above. To determine the tracer $K_D$ in the direct labeling system, 10 nM CoraFluor-1-labeled $His_6$-CRBN/DDB1 complex was added into the assay buffer (25 mM HEPES, 150 mM NaCl, 0.5 mg/mL BSA, 0.005% TWEEN 20, pH 7.5). Thal-FITC was first added in serial dilution (1:2 titration, 9-point, $c_{max}$ = 1 µM) using an HP D300 digital dispenser and allowed to equilibrate for 10 min at room temperature before TR-FRET measurements were taken. The $K_D$ was determined from the one-site model graphed in Prism 10 without any further adjustment.

For ligand displacement assays, 10 nM CoraFluor-1-labeled $His_6$-CRBN/DDB1 complex and 100 nM Thal-FITC were mixed in the assay buffer. Triton X-100 was added to GSS protein stocks to final concentration of 0.1% for ease of dispense. GSS protein samples were added in serial dilution (1:3 titration, 8-point, $c_{max}$ = 3 μM) using an HP D300 digital dispenser and allowed to equilibrate for 10 min at room temperature before TR-FRET measurements were taken. All experiments were performed in technical triplicates. Background signal (bottom) was determined from wells containing 100 μM 5-$NH_2$-lenalidomide to represent complete displacement. The assay ceiling (top) was defined via a no-ligand control. Data were background-subtracted, normalized and fitted to a four-parameter dose-response model [log(inhibitor) vs. response – Variable slope (four parameters)] in Prism 10, with constraints of Top = 1 and Bottom = 0 to derive the ligand $IC_{50}$ values. Ligand $K_D$ values have been calculated using Cheng-Prusoff principles, outlined in the equation below:

$$K_D = \frac{IC_{50}}{1 + \frac{[S]}{K_x}}$$

Where $IC_{50}$ is the measured $IC_{50}$ value, [S] is the concentration of fluorescent tracer, and $K_X$ is the $K_D$ of the fluorescent tracer without adjustment.

For ligand association assays, 10 nM CoraFluor-1-labeled $His_6$-CRBN/DDB1 complex was added to the assay buffer. Triton X-100 was added to GFP proteins stocks to final concentration of 0.1% for ease of dispense. GFP protein samples were added in serial dilution (1:3 titration, 8-point, $c_{max}$ = 1 μM) using an HP D300 digital dispenser and allowed to equilibrate for 10 min at room temperature before TR-FRET measurements were taken. The same serial dilution of each protein sample was performed twice, one without any additive as the sample well and another with 100 μM lenalidomide added as the background well to compete off CRBN-specific association. All experiments were performed in technical triplicates. The corrected TR-FRET ratio was determined by subtracting the ratio of background well from that of the sample well containing the same concentration of GFP sample. Date were fitted to a sigmoidal model [Sigmoidal, 4PL, log(concentration)] in Prism 10, with constraints of Top = 0.4574 (highest mean found for GFP-VAVLDcN) and Bottom = 0 to derive the $K_D$ values.

**General western blotting protocol.** For the tissue culture cells, samples were lysed in Pierce IP lysis buffer supplemented with 1× protease/phosphatase inhibitor cocktail, incubated on ice for 10 min, and clarified by centrifugation (21,000 × g, 4 °C, 10 min). The supernatant of each sample was transferred to a new tube and a BCA assay was performed to measure the protein concentration. The lysates were adjusted to 1.5–2.5 mg/mL, mixed with 5× SDS-PAGE loading buffer (1× concentration: 50 mM Tris–HCl, 2% SDS, 10% glycerol, 1% β-mercaptoethanol, 0.02% bromophenol blue) and boiled at 95 °C for 5 min. For the analysis of recombinant proteins, the protein solution was directly mixed with SDS-PAGE loading buffer and boiled at 95 °C for 5 min. The denatured samples (8–10 μL) were loaded on a 12% Criterion TGX 26-well precast gel or 15-well homemade gel and run at 150 V for 60 min in Tris/Glycine/SDS buffer. The gel was transferred to a nitrocellulose membrane using an Invitrogen iBlot 2 dry blotting system with the preset P0 program. The membrane was cut, blocked with 5% BSA/TBST or 5% milk/TBST and incubated with the primary antibody at 1:1000 dilution at 4°C overnight. The membrane was washed 3× with TBST and incubated with the secondary antibody at 1:10,000 dilution at 24 °C for 1 h, after which it was washed 3× with TBST again. The washed membrane was imaged by IR800 or chemi channel on an Azure imager.

**In vitro ubiquitination of tagged proteins.** 20.0×$10^6$ HEK-CRBN cells per pellet were collected, flash-frozen with liquid nitrogen, and stored at –80 °C until use. Each pellet was lysed in Pierce IP lysis buffer supplemented with 1× protease/phosphatase inhibitor cocktail (1200 µL) and 200 µL of TBS-washed anti-FLAG M2 beads were added to the soluble fraction of lysate. Samples were incubated at 4 °C on a roller for 1 h. Using a magnetic tube rack, the beads were collected and washed 3× with 1 mL TBS. Each sample was then eluted by adding 100 µL of 150 ng/µL 3× FLAG peptide/TBS and incubating at 4 °C on a roller for 1 h. The eluate was collected. Tagged proteins were diluted in PBS, using the same protein concentration for all samples in each experiment. Ubiquitination master mixes were prepared at 2× with and without E1 and E2 enzymes. Final concentrations (1×) of the master mix components were: 0.15 µM UBE1, 2 µM UbcH5a, 1 µM UbcH5c, 400 µg/mL K0 ubiquitin, 1 µM ubiquitin aldehyde, 10 mM Mg-ATP, 1× E3 ligase buffer (50 mM HEPES pH 7.4, 50 mM NaCl, 1 mM TCEP), 10 µM MG132, 100 nM MG101. Reactions were prepared by combining 6.25 µL FLAG eluate, 5.25 µL of substrate (final concentration: 1.26 µM for GSS and 1.05 µM for GFP), 1 µL of 25× DMSO or lenalidomide stock in 2.5% DMSO/PBS (final concentration: 100 µM), and 12.5 µL of ubiquitination master mix in Eppendorf tubes. Reactions were incubated at 37 °C for 60 min then were stopped by the addition of 6.25 µL of 5× SDS-PAGE loading buffer. Samples were heated at 95 °C for 5 min prior to analysis by western blotting.

**Electroporation of GFP-VAVLDcN and GFP-VAVLDN.** HEK293T cells were grown to 80–90% confluency in DMEM+/+ prior to electroporation. Cells were detached by trypsinization as necessary and washed with PBS, then resuspended in PBS and counted. For each sample, an equal number of cells (7.0×$10^5$) were aliquoted into Eppendorf tubes and pelleted. Electroporation mixes were prepared for each sample type. Electroporation mixes were prepared by combining 41.6 µL Neon buffer R containing DMSO or 200 µM lenalidomide, 4.2 µL of 50 µM mCherry protein as the internal standard, and 4.2 µL 50 µM GFP proteins. A mock electroporation mix was also prepared by substituting the mCherry and GFP protein with PBS. Immediately prior to each electroporation, the PBS was removed from the pelleted cells and the pellet was resuspended in 12 µL of electroporation mix. The sample was taken up into a 10 µL tip attached to a Neon pipette, and the pipette tip was submerged in a Neon cuvette containing 3 mL Neon buffer E2. The sample was then electroporated (1150 V, 20 msec, 2 pulses). The cells were then dispensed into 100 µL warmed PBS and flicked to mix. This process was repeated for each sample, with each tip used for 3 electroporation cycles. Cells were then pelleted by centrifugation and the supernatant was removed. Cells were resuspended in 50 µL colorless trypsin-EDTA solution and incubated at 37 °C for 3 min. Trypsinization was quenched by the addition of 0.5 mL colorless DMEM+/+ containing DMSO or 200 µM lenalidomide and the mixture was transferred to a well of a 24-well plate. The samples were then incubated at 37 °C, 5% $CO_2$ until 6 h after electroporation. Each sample was resuspended in 500 µL PBS and analyzed by flow cytometry (PE Texas Red and FITC on FACSymphony A3 Lite). To calculate normalized GFP level, the arithmetic mean of the GFP and mCherry fluorescence intensities for each sample were corrected by subtracting the corresponding intensity values in the mock sample; then, these corrected values were used to calculate a GFP/mCherry fluorescence intensity ratio for each sample. Finally, the resulting values were normalized to the mean ratio for GFP-VAVLDN without lenalidomide.

**Generation of GSS reporter cell lines.** pLenti-GFP-GSS(WT) (plasmid 11) was derived from the plasmids pcDNA3.1-FLAG-GSS (Genscript) and pLenti-GFP-CK1α. The insert corresponding to GSS full-length cDNA and backbone excluding CK1α were amplified using overhang PCR with primers 17–20 and assembled using HiFi DNA Assembly Kit (NEB). The ligated product was transformed into NEB Stable competent cells and the sequences validated by Sanger sequencing. pLenti-GFP-GSS(N470A) was constructed from pLenti-GFP-GSS(WT) using primers 21–22 with QuikChange Lightning kit.

For virus production, HEK293T cells in a 10 cm plate were transfected by pMD2.G (6 μg), psPAX2 (4 μg) and pLenti-GFP-GSS(WT/N470A) (10 μg) using TransIT-Pro reagent (20 μL) following the manufacturer's guidelines. Lentiviruses were collected 48 h after transfection, spun down (500 × g, 5 min, 24 °C) and filtered with a 0.45 μm PES filter. For transduction, HEK293T cells in 1 mL DMEM+/– containing 10 μg/mL polybrene were seeded into 6-well plates containing the virus (500 μL) and incubated for 48 h and then selected by 2 μg/mL puromycin for 3 days. The resulting GFP-GSS reporter cell lines were first sorted by MoFlo Astrios (Beckman Coulter) into three bins according to the FITC signal. The resulting "low" bin of WT and "medium" bin of N470A reporter cell line were selected for further evaluation based on their similar level of GFP-GSS fusion construct detected by western blotting.

**Evaluation of GFP-GSS level upon compound treatment by flow cytometry.** GSS reporter cell lines ($0.05\times10^6$ cells per well) were seeded in 12-well plates in 1 mL DMEM+/+. The cells were stabilized for 1 h and then treated with DMSO or 200 μM Boc-FcN. After 48 h incubation, media was replaced to fresh DMEM+/+ and DMSO or 200 μM Boc-FcN was immediately added. This process was repeated once more, and the cells were incubated for 24 h where they reach approximately 95% confluency (5 days of compound treatment in total). Cells were trypsinized, spun down, and resuspended in 500 μL PBS containing 10 μg/mL propidium iodide to allow for exclusion of dead cells. Cells were analyzed by flow cytometry (PE Texas Red and FITC on FACSymphony A3 Lite). A total of 20,000 events were analyzed for each sample and only the signals of live cells were taken to further analysis. Normalized GFP level was determined by dividing the FITC-A value of each sample with the arithmetic mean of DMSO-treated samples.

**CRBN knockdown in reporter cell lines by siRNA.** CRBN siRNA mix and control siRNA-B were purchased from Santa Cruz Biotechnology. siRNA transfection was performed using similar procedures suggested by the manufacturer. GSS reporter cell lines ($0.2\times10^6$ cells per well) were seeded in 6-well plates in 1 mL DMEM + 10% FBS without antibiotics and incubated for 20 h. For each well, siRNA (18 μL of 10 μM stock) was diluted in 100 μL siRNA transfection medium and 18 μL transfection reagent was diluted in another 100 μL of siRNA transfection medium. The siRNA duplex solution was added to the diluted transfection reagent, incubated for 30 min at 24 °C, and further diluted with 800 μL transfection medium. Media was aspirated from the 6-well plates and the mixture was gently layered onto washed cells. The transfection was allowed to proceed for 7 hours. The cells were supplied with 1 mL DMEM + 20% FBS with 2× antibiotics and incubated for 24 hours. The media was then replaced to fresh DMEM+/+ and cells were further incubated for 96 hours (5 days incubation after transfection). Cells were collected and analyzed by western blotting or flow cytometry as described above.

**Generation of cleaved and full-length GSS.** pET28a-$His_6$-GSS[1–470] (cleaved) or pET28a-$His_6$-GSS[1–474] (full-length) was constructed from plasmid 9 by first replacing the LPETG-$His_6$ tag with residues 464–470 or 464–474 of native GSS sequence using primers 23–24 or 25–26, respectively. A N-terminal $His_6$ tag was then inserted into each plasmid using primers 27–28.

BL21(DE3) cells were transformed with each sequence verified plasmid and used to inoculate overnight cultures of LB + 50 μg/mL kanamycin. The overexpression conditions and Ni-bound HiTrap affinity purification were performed as described in "Overexpression and purification of recombinant GSS-LPETG" using the identical stock solutions, buffers and methods for the two constructs. The concentrations of affinity-purified proteins were determined by measuring the A280 for GSS[1–470] ($\varepsilon$ = 33,015 $M^{-1}$ $cm^{-1}$) and GSS[1–474] ($\varepsilon$ = 34,505 $M^{-1}$ $cm^{-1}$).

**Comparative size-exclusion chromatography analysis of cleaved and full-length GSS.** The affinity-purified GSS proteins were injected on a S75 10/300 GL column, pre-equilibrated and run in PBS (injected amount: 8.0 mg) after one round of freeze-thaw cycle. The chromatograms were exported into raw data points and graphed in Prism 10. The peak areas were integrated using the evaluation tool of UNICORN software. The starting affinity-purified proteins and their combined SEC fractions were adjusted to 1–3 μM final concentration and run on a gel followed by Coomassie blue staining. The combined SEC fractions corresponding to monomeric GSS were aliquoted to Eppendorf tubes (final concentration: 150 μM) and subject to three rounds of freeze-thaw cycles, after which the formation of white precipitates was observed for GSS[1–470].

**PROTEOSTAT staining of cleaved and full-length GSS.** Quantification of GSS aggregation was performed using PROTEOSTAT protein aggregation assay. In brief, affinity-purified proteins were diluted to the desired concentrations by 1× assay buffer. The supplied positive and negative standards were diluted to 20 μM as suggested by the manufacturer. 1× PROTEOSTAT stain (2 μL) was added into each well of a 96-well black wall microplate with clear bottom. Each of the diluted protein or standard (98 μL) was added to each well and the microplate was incubated at 24 °C for 15 min in the dark. Fluorescence measurements were acquired on a SpectraMx iD5 plate reader with SoftMax Pro software version 7.1.2 at 350 nm excitation and 616 nm emission both using filters. Relative fluorescence was calculated by subtracting the fluorescence of the well with assay buffer only from each sample well and normalizing to the corrected fluorescence of negative standard.

**Preparation of insoluble proteins from cell lines for mass spectrometry analysis.** The differential centrifugation method was adapted from Chen and coworkers(7). HEK293T and MOLM-13 cells (55–75×$10^6$ cells per sample) were collected in Falcon tubes. 1 mL lysis buffer (30 mM Tris-HCl, 40 mM NaCl, 1 mM DTT, 3 mM $CaCl_2$, 3 mM $MgCl_2$, 5 % glycerol, 1 % triton X-100, EDTA-free protease inhibitor, pH 7.5) per 50×$10^6$ cells was added to each pellet. Samples were pipetted vigorously up and down and chilled on ice for 15 min. 1 mL of each suspension was transferred to an Eppendorf tube and spun down at 800 × g, 10 min at 4 °C to remove the cell debris. The supernatant was transferred to a clean tube and treated with 1 μg/mL RNase and 20 units/mL DNase for 30 min on ice to exclude other insoluble nucleic acids and proteins that aggregate as a result of nucleic acid binding. The resulting solution was further spun down at 10,000 × g, 15 min at 4 °C and the supernatant was labeled as [WCL]. The insoluble pellet was taken to be solubilized by 5% SDS, 50 mM TEAB pH 7.55 and spun down at 21,000 × g, 15 min at 4 °C, which yielded in the Triton-insoluble fraction and a small pellet that was insoluble in SDS. The concentrations of WCL, Triton-insoluble fraction, and SDS-insoluble fraction (lysed in 8 M urea/PBS) were initially determined by reductant-compatible BCA kit (ThermoFisher 23250). The SDS-insoluble pellet contained minimal protein whereas WCL and Triton-insoluble fractions contained approximately 3 mg and 50 μg protein, respectively. Thus, we labeled the Triton-insoluble fractions as [INSOL] and took [WCL] and [INSOL] samples for further analysis. For mass spectrometry sample preparation, WCL was diluted to the same protein amount as INSOL with 5% SDS, 50 mM TEAB, pH 7.55. Then the WCL and INSOL samples were reduced, alkylated, and digested by trypsin on S-trap as described in "Formation study of C-terminal cyclic imide and its hydrolysis products in recombinant proteins". For comparison of WT and CRBN-KO samples, 4 technical replicates were generated by dividing WCL and INSOL samples into 4 equal volumes before reduction by dithiothreitol. After the S-trap digestion, each dried replicate was resuspended in 10 μL dd$H_2O$, labeled with TMT 10-plex reagent (10 μL) at 24 °C for 1 h, and quenched with 1 M Tris-Cl pH 7.6 (5 μL) for 15 min. The samples were combined and dried on a vacufuge. The resulting samples were fractionated into 4 fractions using the Pierce high pH reversed-phase peptide fractionation kit. The peptides were eluted sequentially by 5% (excluded from analysis), 10%, 15%, 25% and 50% acetonitrile/0.1% TEA. Immediately after the elution, each fraction was acidified by the addition of 5 μL of 10% formic acid. The dried fractions were resuspended in 20 μL of 0.1% formic acid and injected on an Orbitrap Eclipse Tribrid Mass Spectrometer coupled with a Vanquish Neo HPLC system using the same method as "Proteomics mass spectrometry acquisition procedures". Analysis was performed in Thermo Scientific Proteome Discoverer version 2.4.1.15. The raw data were searched against SwissProt human (Homo sapiens) protein database (21 February 2019; 20,355 total entries) and contaminant proteins using the Sequest HT algorithm. Searches were performed with the following guidelines: spectra with a signal-to-noise ratio greater than 1.5; mass tolerance of 20 ppm for the precursor ions and 0.02 Da for the fragment ions; semi-trysin digestion at the peptide N-terminus only; fewer than 2 missed cleavages; static carboxyamidomethylation of cysteine residues (+57.021 Da); static TMT 10-plex labeling (+226.163 Da) at lysine residues and N-termini only for the TMT-labeled samples; variable oxidation on methionine residues (+15.995 Da); variable dehydration on asparagine and glutamine residues at C-terminus only (–18.015 Da). The TMT reporter ions were quantified using the Reporter Ions Quantifier node and normalized to the total peptide amount. PSMs were filtered using a 1% false discovery rate (FDR) using Target Decoy PSM Validator. For the quantification of total proteins, the data were further filtered to include only master proteins with high protein FDR confidence, at least 3 unique peptides, and exclude all contaminant proteins. To derive the list of peptides generated from internal cleavage at N or Q, the data were filtered to include only peptides without oxidation modifications, bearing N or Q residues at the peptide C-terminus with or without dehydration, and derived from non-contaminant proteins. The peptides that were mapped back to the protein C-terminus were excluded. The abundance ratios and their associated p-values for the TMT-labeled samples were calculated by one-way ANOVA with TukeyHSD post-hoc test.

#### III. Materials and Instrumentation

General Supplies

- (*R*, *S*)-Lenalidomide (BioVision 1862-25)
- MLN4924 (Selleck Chemicals S7109)
- Zeba 7 kDa desalting columns (0.5 mL and 5 mL, ThermoFisher 89882 and 89892)
- Dulbecco's Modified Eagle's Medium (DMEM) (Genesee Scientific 25-500)
- Trypsin-EDTA (Fisher Scientific 25200114)
- Fetal bovine serum (FBS) (Peak Serum PS-FB2)
- Penicillin-streptomycin (100×) (Lonza 17-602E)
- Pierce IP lysis buffer (ThermoFisher 87788)
- Protease/phosphatase inhibitor cocktail (100×) (Cell Signaling Technology 5872S)
- Protease inhibitor cocktail (Sigma-Aldrich 11873580001, 1 tablet dissolved in 2 mL water for a 25× stock solution)
- BCA solution (BCA Reagent A) (VWR 786-847)
- Copper Solution (BCA Reagent B) (VWR 76825-860)
- 12% Criterion TGX precast gels (Bio-Rad 5671044)
- iBlot 2 nitrocellulose transfer stack (Invitrogen IB23001; IB23002)
- Vivaspin spin concentrators (Cytiva)
- Opti-MEM I Reduced Serum Medium (ThermoFisher 31985070)
- TransIT-Pro Transfection Reagent (Mirus MIR 5760)

Cloning Reagents

- Q5 site-directed mutagenesis kit (New England BioLabs E0552S)
- QuikChange Lightning Site-directed Mutagenesis Kit (Agilent 210518)
- HiFi DNA Assembly Kit (New England BioLabs E5520S)

Mass Spectrometry

- S-Trap micro (Protifi)
- Triethylammonium bicarbonate buffer (Sigma-Aldrich T7408-100ML)
- Pierce high pH reversed-phase peptide fractionation kit (ThermoFisher 84868)
- LysC (Promega, VA1170)
- Trypsin (Promega, VA5117)
- TMTpro 16-plex (ThermoFisher, A44520)
- TMT 10-plex (ThermoFisher, 90406)
- Pierce Peptide Desalting Spin Column (ThermoFisher, 89852)
- 96-well plate, 1.0 mL, round wells (Agilent, 5043-9305)

TR-FRET

- Proxiplate-384 Plus (VWR PERK6008280)

In Vitro Ubiquitination

- Anti-FLAG M2 beads (Sigma-Aldrich M8823-1ML)
- 3× FLAG peptide (Sigma-Aldrich F4799-4MG)
- UBE1 (R and D Systems E-305-025)

- UbcH5a (R and D Systems E2-616-100)
- UbcH5c (R and D Systems E2-627-100)
- K0 ubiquitin (R and D Systems UM-NOK-01M)
- Ubiquitin aldehyde (SCBT sc-4316)
- Mg-ATP (R and D Systems B-20)
- MG132 (Selleck Chemicals S2619)
- MG101 (Tocris 3358)

Electroporation
- Neon Transfection System 10 µL Kit (ThermoFisher MPK1096)
- Neon Transfection Tubes (ThermoFisher MPT100)
- DMEM with 4.5 g/L Glucose, without phenol red (Lonza, 12-917F)
- Trypsin-EDTA (0.5%), no phenol red (ThermoFisher, 15400054)

siRNA knockdown
- siRNA transfection medium (SCBT sc-36868)
- siRNA transfection reagent (SCBT sc-29528)
- Control siRNA-B (SCBT sc-44230)
- CRBN siRNA (SCBT sc-78528)

Aggregation analysis
- PROTEOSTAT protein aggregation assay (Enzo Life Sciences ENZ-51023)
- 96-well black/clear bottom plate (ThermoFisher 165305)
- RNase A (ThermoFisher EN0531)
- DNase (New England BioLabs M0303S)
- Reductant-compatible BCA kit (ThermoFisher 23250)

Recombinant Proteins
- Recombinant Glutathione Synthetase full-length (Novus Biologicals NBP1-50862)

Mammalian Cell Lines
HEK293T cells were obtained from American Type Culture Collection (ATCC). HEK293FT cells stably expressing FLAG-CRBN (HEK-CRBN cells) were kindly provided by the Deshaies Lab (California Institute of Technology). MOLM-13 cells were kindly provided by the Kharas Lab (Memorial Sloan Kettering Cancer Center).

Bacterial Strains
*E. coli* 5-alpha Competent (High Efficiency) (New England BioLabs C2987H)
*E. coli* BL21 (DE3) (New England Biolabs C2527H)
*E. coli* NEB Stable Competent (New England BioLabs C3040I)

Purchased Synthetic Peptides
- VEALGNESALE (Genscript)
- VEALANESALE (Genscript)
- VEALKNESALE (Genscript)

- VEALPNESALE (Genscript)
- VEALFNESALE (Genscript)
- VEALLQPKALE (Genscript)
- VEALLQERALE (Genscript)
- VEALLQESALE (Genscript)
- HBB[42−60]: FFESFGDLSTPDAVMGNPK (Biomatik)
- ACTB[96−113]: VAPEEHPVLLTEAPLNPK (Biomatik)
- ACTB[247–257]: VITIGNERFRC (Genscript)
- CALM3[49–59]: LQDMINEVDAD (Genscript)
- CPSM[460–470]: VLMNPNIASVQ (Genscript)
- HBA[74–84]: VDDMPNALSAL (Genscript)
- GFAP[349–359]: YQDLLNVKLAL (Genscript)
- ACTB[105–111]: LTEAPLN (Genscript)
- ACTB[246–252]: QVITIGN (Genscript)
- CALM3[48–54]: ELQDMIN (Genscript)
- GSS[453–474]: AIEHADGGVAAGVAVLDNPYPV (Genscript)
- GSS[464–470]: GVAVLDN (Genscript)
- GSS sorttag-N: GGGVAVLDN (Genscript)

<u>Antibodies</u>

| ***No.*** | ***Antibody Name*** | ***Host Species*** | ***Blocking Buffer*** | ***Supplier*** | ***Catalog#*** |
|---|---|---|---|---|---|
| 1 | $His_6$ | Mouse mAb | For TR-FRET | Abcam | ab18184 |
| 2 | GSS | Rabbit pAb | 5% milk/TBST | Proteintech | 15712-1-AP |
| 3 | GFP (D5.1) | Rabbit mAb | 5% BSA/TBST | Cell Signaling Technology | 2956S |
| 4 | FLAG | Rabbit mAb | 5% BSA/TBST | Cell Signaling Technology | 14793S |
| 5 | CRBN | Rabbit mAb | 5% BSA/TBST | Cell Signaling Technology | 71810 |
| 6 | β-actin (C4) | Mouse mAb | 5% BSA/TBST | Santa Cruz Biotechnology | sc-47778 |
| 7 | Vinculin (V284) | Mouse mAb | 5% milk/TBST | Bio-Rad | MCA465 GA |
| 8 | Anti-mouse-HRP | Goat pAb | Secondary antibody | Rockland Immunochemicals | 610-1302 |
| 9 | Anti-rabbit-HRP | Goat pAb | Secondary antibody | Rockland Immunochemicals | 611-1302 |
| 10 | Anti-mouse-IRDye® 800CW | Goat pAb | Secondary antibody | LI-COR Biosciences | 925-32210 |

<u>Plasmids</u>

| ***No.*** | ***Plasmid name*** | ***Source*** |
|---|---|---|
| 1 | pET28a-GFP-LPETG-$His_6$ | Previous work(1) |
| 2 | pET28a-GFP-LPETGGGFNPK-$His_6$ | This work |
| 3 | pET28a-GFP-LPETGGGFNAK-$His_6$ | This work |
| 4 | pET28a-GFP-LPETGGGFNLK-$His_6$ | This work |
| 5 | pET28a-GFP-LPETGGGFNEK-$His_6$ | This work |
| 6 | pET28a-GFP-LPETGGGFNEEK-$His_6$ | This work |
| 7 | pET28a-GFP-LPETGGGFNEVK-$His_6$ | This work |
| 8 | pcDNA3.1-FLAG-GSS | Genscript |
| 9 | pET28a-GSS[1–463]-LPETG-$His_6$ | This work |
| 10 | pLenti-GFP-CK1α-puro | Previous work(8) |
| 11 | pLenti-GFP-GSS(WT)-puro | This work |
| 12 | pLenti-GFP-GSS(N470A)-puro | This work |
| 13 | pET28a-$His_6$-GSS[1–470] | This work |
| 14 | pET28a-$His_6$-GSS[1–474] | This work |

<u>Primers</u>

| ***No.*** | ***Primer name*** | ***Sequence (5' to 3')*** |
|---|---|---|
| 1 | GGFNPK-F | AATCCGAAACACCACCACCACCACCAC |
| 2 | GGFNPK-R | AAAGCCACCACCGGTTTCCGGGAGGCT |
| 3 | NAK-F | TGGTGGTGTTTCGCATTAAAGCCACCACCGG |
| 4 | NAK-R | CCGGTGGTGGCTTTAATGCGAAACACCACCA |
| 5 | NLK-F | GTGGTGGTGGTGTTTCAGATTAAAGCCACCACC |
| 6 | NLK-R | GGTGGTGGCTTTAATCTGAAACACCACCACCAC |
| 7 | NEK-F | TGGTGGTGGTGTTTCTCATTAAAGCCACCACCGGTTTC |
| 8 | NEK-R | GAAACCGGTGGTGGCTTTAATGAGAAACACCACCACCA |
| 9 | NEEK-F | TGGCTTTAATGAAGAAAAACACCACCACCACCACCACTGAATG |
| 10 | NEEK-R | CCACCGGTTTCCGGGAGG |
| 11 | NEVK-F | TGGTGGCTTTAATGAAGTAAAACACCACCACCACC |
| 12 | NEVK-R | GGTGGTGGTGGTGTTTTACTTCATTAAAGCCACCA |
| 13 | pET28a-LPETG-F | CTCCCGGAAACCGGTCAC |

| | | |
|---|---|---|
| 14 | pET28a-LPETG-R | GGTATATCTCCTTCTTAAAGTTAAACAAAATTATTTCTAGAG |
| 15 | GSS[1–463]-F | CTTTAAGAAGGAGATATACCATGGCCACCAACTGGGGG |
| 16 | GSS[1–463]-R | TGGTGACCGGTTTCCGGGAGCGCTGCCACACCACCATC |
| 17 | pLenti-GFP-F | TAATAGCTGCAGATATCCAGCACAGTG |
| 18 | pLenti-GFP-R | GGTGGCGGAGAGTCCGGAC |
| 19 | GSS(WT)-F | AGTCCGGACTCTCCGCCACCATGGCCACCAACTGGGGG |
| 20 | GSS(WT)-R | CTGGATATCTGCAGCTATTACACAGGGTATGGGTTGTCCAG |
| 21 | GSS(470A)-F | AGTCCTGGACGCCCCATACCCTGTGTAATAGCTG |
| 22 | GSS(470A)-R | GCCACTCCCGCTGCC |
| 23 | Replace-470-F | TTCTGGATAACTGAGATCCGGCTGCTAAC |
| 24 | Replace-470-R | CCGCAACACCCGCTGCCACACCACC |
| 25 | Replace-474-F | ATAACCCGTATCCGGTGTGAGATCCGGCTGCTAAC |
| 26 | Replace-474-R | CCAGAACCGCAACACCCGCTGCCACACCACC |
| 27 | NterHis-F | CACCACCACGCCACCAACTGGGGGAGC |
| 28 | NterHis-R | GTGGTGGTGCATGGTATATCTCCTTCTTAAAGTTAAACAAAATTATTTCTAGAGG |

Instrumentation

Protein quantification by bicinchoninic acid assay (BCA) and TR-FRET measurements were performed on multi-mode microplate reader SpectraMax iD5 (Molecular Devices LLC). Protein concentration and OD600 measurements were measured by Nanodrop One$^{C}$ Microvolume UV-Vis Spectrophotometer (ThermoFisher). Cell lysis was performed using Branson Ultrasonic Probe Sonicator (model 250). Fluorescence and chemiluminescence imaging were performed using Azure Imager c600 or 400 (Azure Biosystems, Inc., Dublin, CA). Protein purification and analytical SEC was performed using an ÄKTA pure 25 equipped with F9-R fraction collector, C9n conductivity monitor, and computer running UNICORN v6.3.2.89 (GE Healthcare). Reverse-phased HPLC for peptide fractionation and protein purification was performed using an Agilent 1260 Infinity II system. All proteomics data were obtained on Vanquish Neo HPLC system (ThermoFisher) connected in line to Orbitrap Fusion Lumos Tribrid Mass Spectrometer or Orbitrap Eclipse Tribrid Mass Spectrometer (both ThermoFisher) within the Mass Spectrometry and Proteomics Resource Laboratory at Harvard University. Intact protein mass spectra were collected using Bruker Impact II q-TOF mass spectrometer coupled to Agilent 1290 HPLC within the Mass Spectrometry and Proteomics Resource Laboratory at Harvard University. Western blotting transfer was performed using Invitrogen iBlot 2 dry blotting system. Electroporation was performed using Neon electroporation system (ThermoFisher). Flow cytometry was conducted using FACSymphony A3 Lite analyzer (BD) and cell sorting was conducted using MoFlo Astrios (Beckman Coulter). Cell numbers and viability were measured using TC20 automated cell counter (Bio-Rad). Samples were dried using Vacufuge Plus (Eppendorf).

Software

Data was analyzed and visualized using Microsoft Excel (v16.44) and GraphPad Prism (v8.4.3). DNA and protein sequences were analyzed using Geneious (v11.0.3). Proteomics data was analyzed using Xcalibur Qual Browser (v3.0.63) and Proteome Discoverer (v2.4.1.15). Flow cytometry populations were distinguished using BD FACSDiva (v8.0.1). Images were made using ImageJ (NIH, v1.52q), Adobe Photoshop (v21.1.1) and Adobe Illustrator (v24.1).

#### IV. Synthetic Procedures

General Procedure for Fmoc Solid-Phase Peptide Synthesis

Peptides were synthesized by Fmoc-SPPS on 2-chlorotrityl chloride (2-CTC) resin (loading 1.0–2.0 mmol/g, 0.25 mmol scale). 2-CTC resin was swelled in DMF for 1 h. The first Fmoc-amino acid (4 equiv) activated with DIPEA (8 equiv) in DMF was doubly coupled to the 2-CTC resin (loading 1.0–2.0 mmol/g, 0.25 mmol scale) for 30 min. Fmoc-deprotection was carried out with 20% piperidine in DMF (10 min × 2). Fmoc-amino acids (1 mmol in 5 mL of DMF, 4 equiv) were activated with HATU (1 mmol in 5 mL of DMF, 4 equiv) and DIPEA (2 mmol in 5 mL of DMF, 8 equiv) for 5 min and allowed to couple for 30 min with constant shaking. Boc-protected amino acids were employed as last amino acids for the synthesis of global protected peptides. The resulting resins were washed with DMF (× 3) and DCM (× 3), and dried under vacuum.

General Procedure for Cleavage from the Resin

The peptide was cleaved using Hexafluoroisopropanol (HFIP)/DCM (1:4 v/v) cocktail for 30 min. The cleavage mixture was filtered, and the resin was washed with DCM. The combined solutions were evaporated by $N_2$ bubbling to minimum volume, and the crude peptides were dissolved in MeCN:water (1:1) and further diluted to around 25% MeCN with water and lyophilized. The crude peptides were used for next step without purification.

General Procedure for the Synthesis of Target Peptides-cN

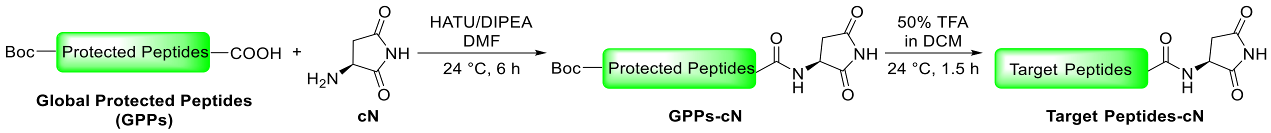


Crude **GPPs** (0.100 mmol, 1.00 equiv) and **cN** (0.30 mmol, 3.00 equiv) were dissolved in DMF (2.0 mL, 0.050 M). DIPEA (0.600 mmol, 6.00 equiv) and HATU (0.3 mmol, 3.00 equiv) were added sequentially to the stirred reaction mixture. After stirring at 24 °C for 6 h, DMF was removed with the aid of a rotary evaporator.

The crude material was dissolved in 2 mL of 50% TFA solution (in DCM, with 3% triisopropylsilane (TIPS), additional 3% ethanedithiol was used for the peptides containing Met). After stirring at 24 °C for 1.5 h, the obtained mixture was concentrated by $N_2$ bubbling to minimum volume, after which 1 mL of DMF was added. The mixture was purified by semi-preparative HPLC to yield **Target peptides-cN**.

High Performance Liquid Chromatography (HPLC) and mass spectrometry (MS)

Analytical reversed-phase (RP) HPLC analyses were performed on an Agilent 1260 Infinity II HPLC with UV detection (220 nm and 256 nm) using a XB-C18 column (3.6 µm, 4.6 × 250 mm) and an Agilent Prep-C18 column (5 µm, 30 ×100 mm). Linear gradients of MeCN (with 0.1 % TFA, buffer B) in water (with 0.1 % TFA, buffer A) were used for all systems to elute bound peptides. The flow rates were 1 mL/min (analytical) and 20 mL/min (C18 preparative).

Low-resolution mass spectrometry (LRMS) measurements were obtained on a Waters Acquity UPLC equipped with SQ Detector 2 mass spectrometer.

#### V. Catalog of LC-MS traces

**VEALLcN.** UPLC-MS (BEH-C18 column: 1.7 µm, 130Å, 2.1 × 150 mm at 220 nm) of LTEAPLcN (obs. 640.4330; calc. for $[M + H]^+$ 640.3665).

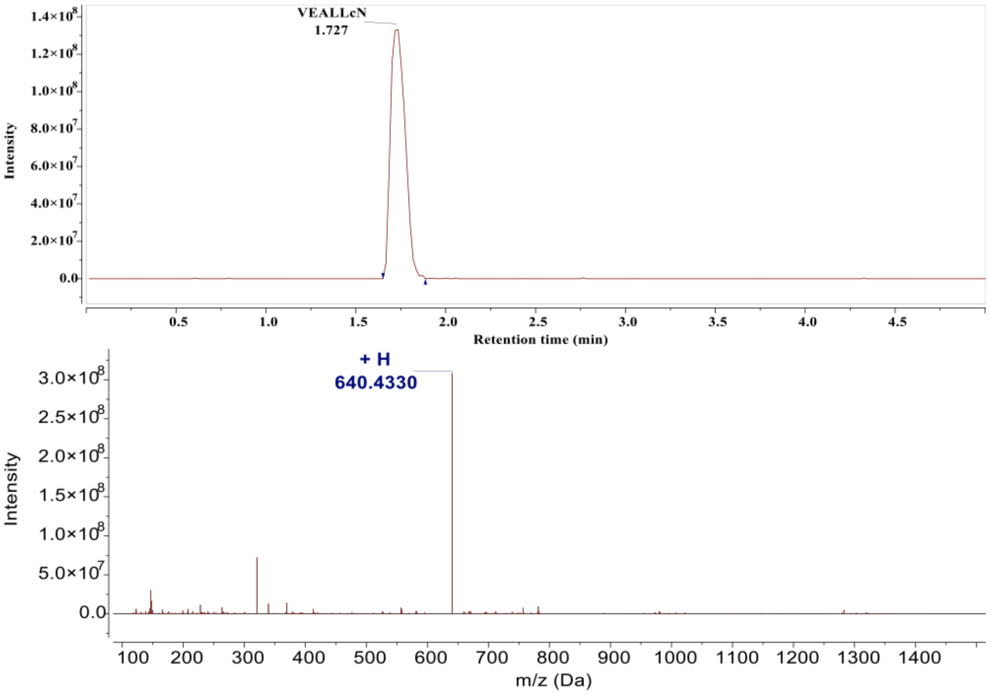


**LTEAPLcN**. Analytical HPLC (XB-C18 column: 3.6 µm, 4.6 × 250 mm at 220 nm) and ESI-MS of LTEAPLcN (obs. 739.8298, 761.3965; calc. for $[M + H]^+$ 739.8480, $[M + Na]^+$ 761.3804).

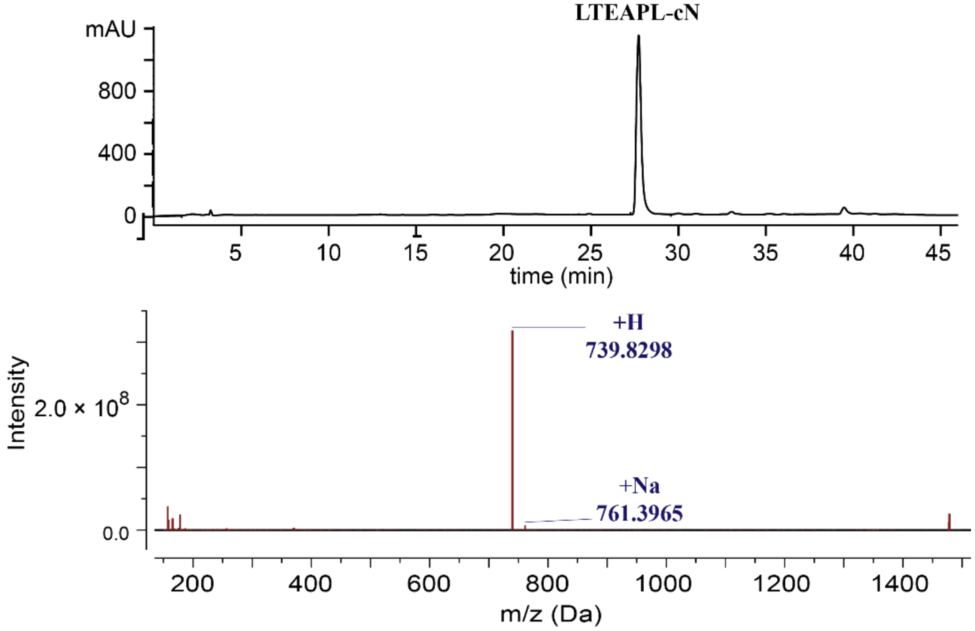

**ELQDMIcN**. UPLC-MS (BEH-C18 column: 1.7 μm, 130Å, 2.1 × 150 mm at 220 nm) of ELQDMIcN (obs. 844.7949, 866.3616; calc. for $[M + H]^+$ 844.9590, $[M + Na]^+$ 866.3694).

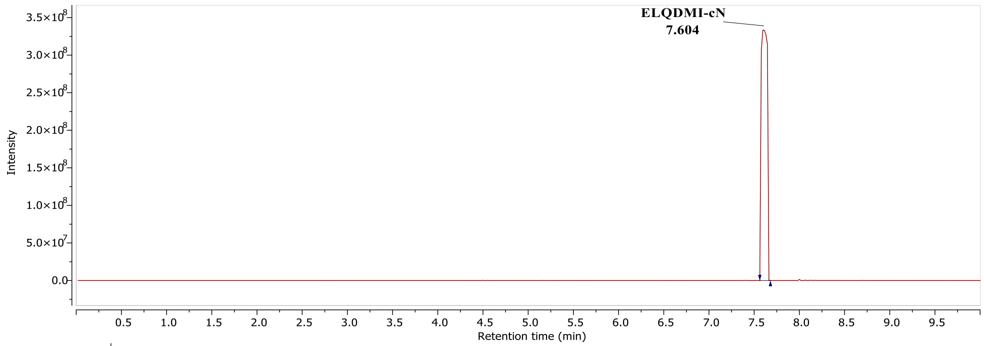


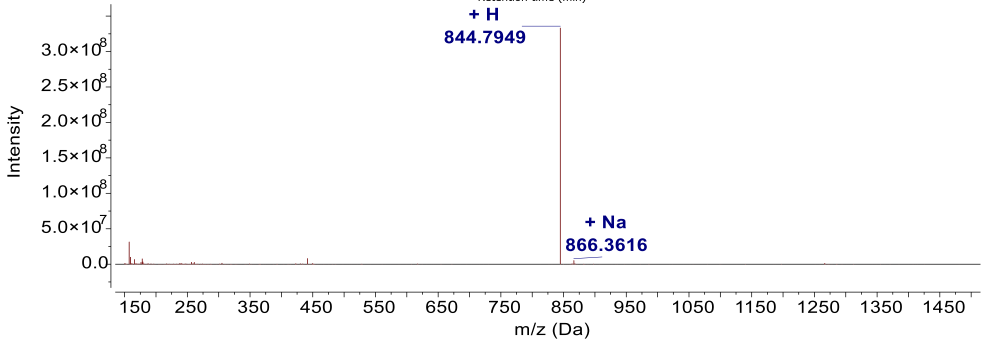


**QVITIGcN.** UPLC-MS (BEH-C18 column: 1.7 μm, 130Å, 2.1 × 150 mm at 220 nm) of QVITIGcN (obs. 726.4307, 748.4362; calc. for $[M + H]^+$ 726.4145, $[M + Na]^+$ 748.3964).

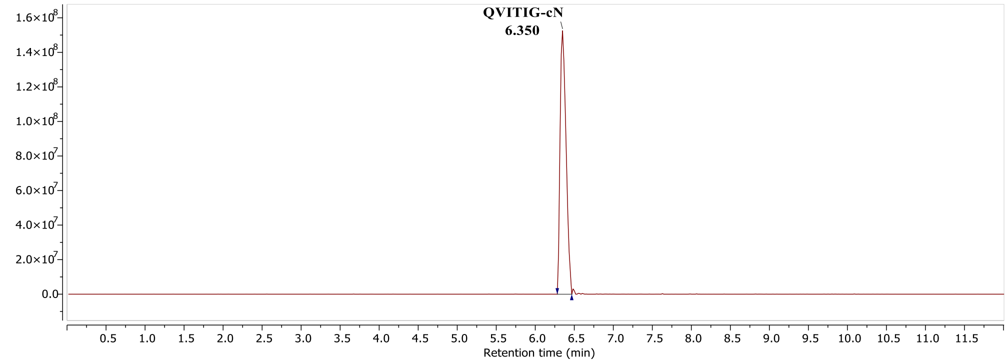


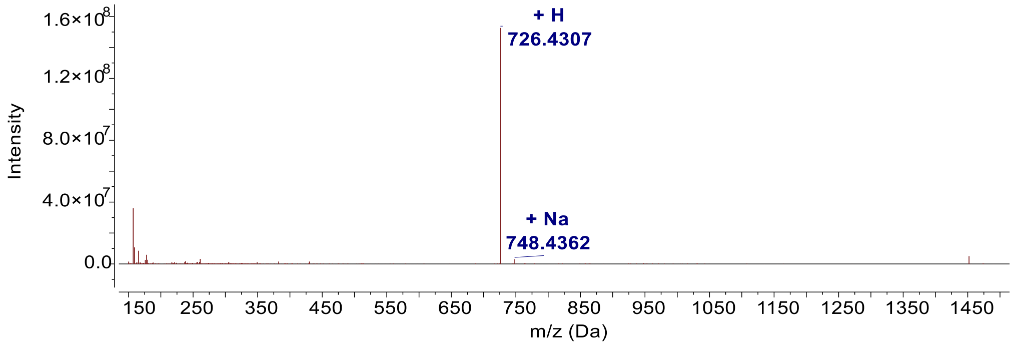

**GVAVLDcN.** UPLC-MS (BEH-C18 column: 1.7 µm, 130Å, 2.1 × 150 mm at 220 nm) of GVAVLDcN (obs. 669.3483, 691.2932; calc. for $[M + H]^+$ 669.3566, $[M + Na]^+$ 691.3386).

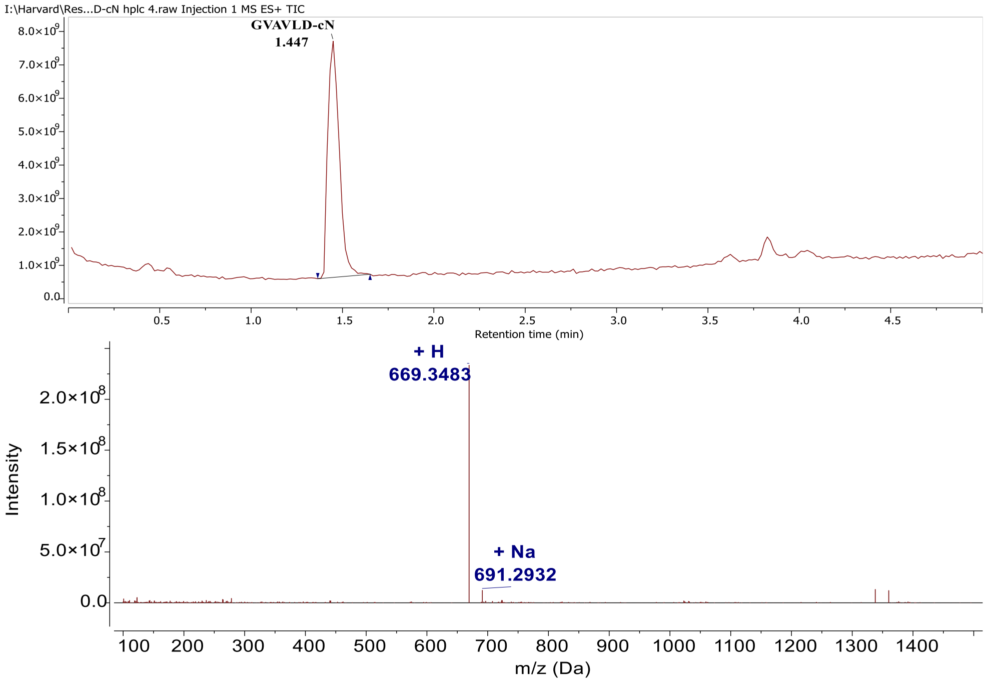


**GGGVAVLDcN.** UPLC-MS (BEH-C18 column: 1.7 µm, 130Å, 2.1 × 150 mm at 220 nm) of GGGVAVLDcN (obs. 783.3988, 805.4431; calc. for $[M + H]^+$ 783.3995, $[M + Na]^+$ 805.3815).

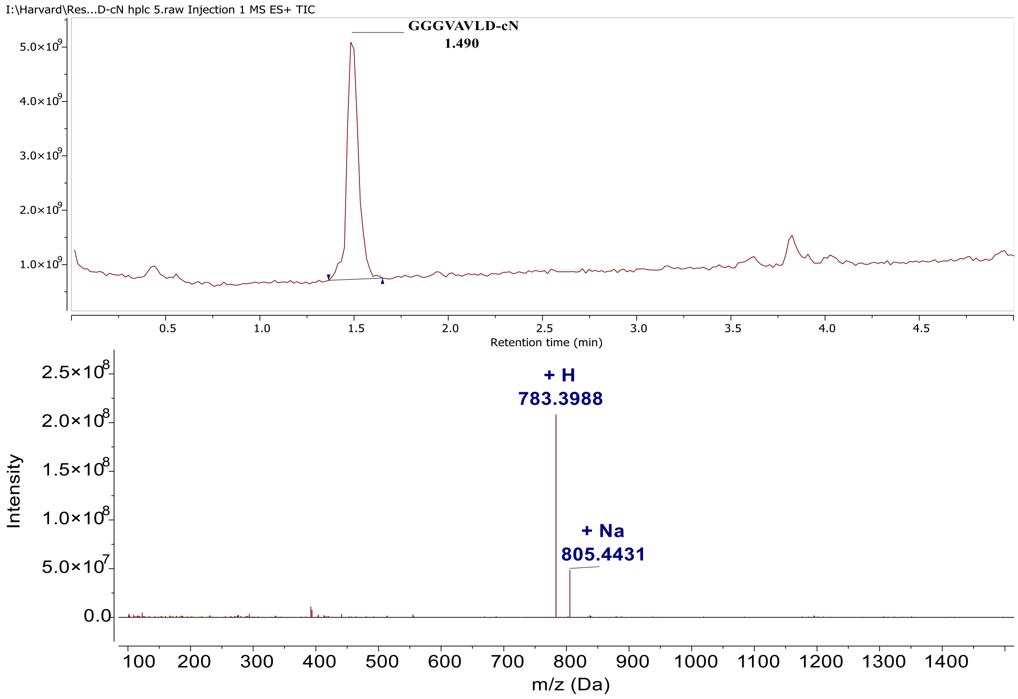

**VI. Legends for Datasets S1–S9**

Dataset S1. Meta-analysis of C-terminal cyclic imide sites from CPTAC datasets.

Dataset S2. Meta-analysis of deamidated sites from CPTAC datasets.

Dataset S3. Results of formation studies shown in Figures 2 and S1.

Dataset S4. Results of formation studies shown in Figures 3 and S2.

Dataset S5. Computational analysis of secondary structural features shown in Figures 4 and S3.

Dataset S6. Results of formation studies and quantitative proteomics shown in Figures 5 and S4.

Dataset S7. Results of formation, binding, and cellular studies shown in Figures 4, 6 and S5.

Dataset S8. Results of aggregation assay and label-free proteomics shown in Figures 7 and S7.

Dataset S9. Results of quantitative proteomics shown in Figures 7 and S7.
